# Nanoparticle-Supported, Rapid, Digital Quantification of Neutralizing Antibodies Against SARS-CoV-2 Variants

**DOI:** 10.1101/2024.11.05.622148

**Authors:** Seyedsina Mirjalili, Md Ashif Ikbal, Ching-Wen Hou, Maziyar Kalateh Mohammadi, Yeji Choi, Laimonas Kelbauskas, Laura A. VanBlargan, Brenda G. Hogue, Vel Murugan, Michael S. Diamond, Chao Wang

## Abstract

The measurement of neutralizing immune responses to viral infection is essential, given the heterogeneity of human immunity and the emergence of new virus strains. However, neutralizing antibody (nAb) assays often require high-level biosafety containment, sophisticated instrumentation, and long detection times. Here, as a proof-of-principle, we designed a nanoparticle-supported, rapid, electronic detection (NasRED) assay to assess the neutralizing potency of monoclonal antibodies (mAbs) against SARS-CoV-2. The gold nanoparticles (AuNPs) coated with human angiotensin-converting enzyme 2 (ACE2) protein as nAb potency reporters were mixed with the mAbs to be tested, as well as streptavidin-conjugated multivalent spike (S) protein or their receptor binding domains (RBD). High-affinity and ACE2-competitive nAbs alter the S (or RBD)-to-ACE2 binding level and modulate AuNP cluster formation and precipitation. The amount of free-floating AuNP reporters is quantified by a semiconductor-based readout system that measures the AuNPs’ optical extinction, producing nAb signals that can differentiate SARS-CoV-2 variants (Wuhan-Hu-1, Gamma, and Omicron). The modular design nature, short assay time (less than 30 minutes), and portable and inexpensive readout system make this NasRED-nAb assay applicable to measuring vaccine potency, immune responses to infection, and the efficacy of antibody-based therapies.

---

Infectious diseases, such as coronavirus disease 2019 (COVID-19), caused devastatingly high levels of morbidity and mortality with substantial economic losses worldwide^1^. During the COVID-19 pandemic, anti-SARS-CoV-2 monoclonal antibodies (mAbs) were authorized to treat patients with mild to moderate symptoms. These mAbs targeted the spike (S) protein of SARS-CoV-2 virions, which enables cell entry of the virus through binding to the human angiotensin-converting enzyme 2 (ACE2) receptor^2^. Such mAbs are neutralizing antibodies (nAbs) produced by the human immune response^3,4^ or rationally engineered^5–7^ and could block the S protein binding to ACE2 to inhibit viral entry into cells. The S protein and its receptor binding domain (RBD) are the primary antigenic targets of nAbs in diagnostics and vaccine development^8^. The S protein and RBD are under significant selective pressure to mutate to evade host immunity, as exemplified by the rapid and continued emergence of SARS-CoV-2 variants^9,10^. For example, Omicron variants (e.g., XBB.1.5, EG.5.1, JN.1, and KP.3) continue to accrue mutations that confer resistance to immunity^9^, eventually making all prior mAb treatment options ineffective. To improve pandemic preparedness and evaluate treatment resistance for future infections, it is crucial to establish a modular method to identify the quality of the nAbs as the viruses mutate^7,11^.

Commercially available serological assays such as anti-spike enzyme-linked immunosorbent assay (ELISA) focus on quantifying the total Ab levels as a predictor of immunity but do not provide insight into nAb potency^12–14^. Currently, the plaque reduction neutralization test (PRNT) is the gold standard for analyzing nAbs against many viruses, but it is labor-intensive and can require high-level containment facilities that are available only in specialized laboratories^13^; microneutralization assays (MNA) have improved speed but still require several days to generate results, and similar to PRNT, require specialized containment, at least for some emerging viruses. Pseudoviruses offer a safer alternative but face challenges, one of which is that the spike proteins displayed on pseudoviruses may not be identical to those on the authentic virus, potentially affecting the accuracy of the results^13^. Surrogate ELISA assays^12^ and competitive chemiluminescence immunoassay^15^ are promising alternatives in lower biosafety settings but require multi-day workflows and specialized equipment for readout. In comparison, the Lateral Flow Assay (LFA)^16^, emerging microfluidic systems^17^ or electrochemical biodevices^18^ offer higher throughput but suffer from poor sensitivity, require complex enzymatic reactions, or face challenges in achieving rapid and accurate readout and data interpretation. Therefore, there is still a need for low-cost, rapid, facile assays that can effectively assess the neutralization capabilities of antibodies and serum samples against emerging virus variants. Here, we report the development of a low-cost, rapid, sensitive, digital neutralizing assay that can distinguish the nAb potency against different SARS-CoV-2 variants, quantify the neutralizing ability of human sera samples, and report the results in a point-of-care (POC) setting using an inexpensive semiconductor-based, portable electronic detection (PED) system.

## <u>Na</u>noparticle-<u>s</u>upported, <u>r</u>apid <u>e</u>lectronic <u>d</u>etection (NasRED) platform for antibody sensing

NasRED is an in-solution protein sensing platform that aims to minimize the analytical time (<30 minutes), sample volume (<10 µL), cost (reagent cost estimated <$3 per test), and complexity of operation^19–21^. The whole sensing process (**Fig. 1**) is completed in a single tube without washing, labeling, or fluorescent imaging steps but achieved by simply mixing proteins or biological media to be tested with a prepared sensing solution comprising functionalized gold nanoparticles (AuNPs, **Fig. 1a-b**). Fundamentally different from conventional protein assays, such as ELISA or LFA, which require a stationary surface and passive reagent diffusion for reaction and readout, NasRED utilizes centrifugal forces to both enhance the active diffusion of proteins and the reagent-bound AuNPs and concentrate reagents at the bottom of the tube (**Fig. 1c**). This approach shortens the assay time by minimizing the precipitation length of AuNPs and promotes protein detection at a low reagent concentration, exemplified by sub-femtomolar SARS-CoV-2 nucleocapsid (N) -protein detection in various biological fluids^21^. Following a brief (*e.g.*, ∼20 min or shorter) incubation (**Fig. 1d**), a vortex agitation is introduced to preserve the specificity of the reaction by returning unbound AuNPs back into the solution without breaking up the biochemically formed AuNP clusters (**Fig. 1e**). Therefore, the protein binding signals are transduced to AuNP aggregation and precipitation at the bottom of the tube, subsequently to plasmonic absorption modulation of free-floating AuNPs in the supernatant, and finally to electronic circuit signals. The signal digitization is accomplished by a semiconductor-based PED system (**Fig. 1f**). The signal-collecting module, made of a light emitting diode (LED), a photodetector (PD), and a microcentrifuge tube chamber to block ambient light, measures the AuNP concentration within the optical path between the LED and PD and converts the optical extinction of the AuNPs to electronic current signals. The signal processing module is built onto a circuit board that stabilizes the LED and PD signals for reliable protein detection and processes the signals for external communications using Wi-Fi, USB, or Bluetooth connections (**Fig. 1g and Fig. S1**)^21^.

**Figure 1.**
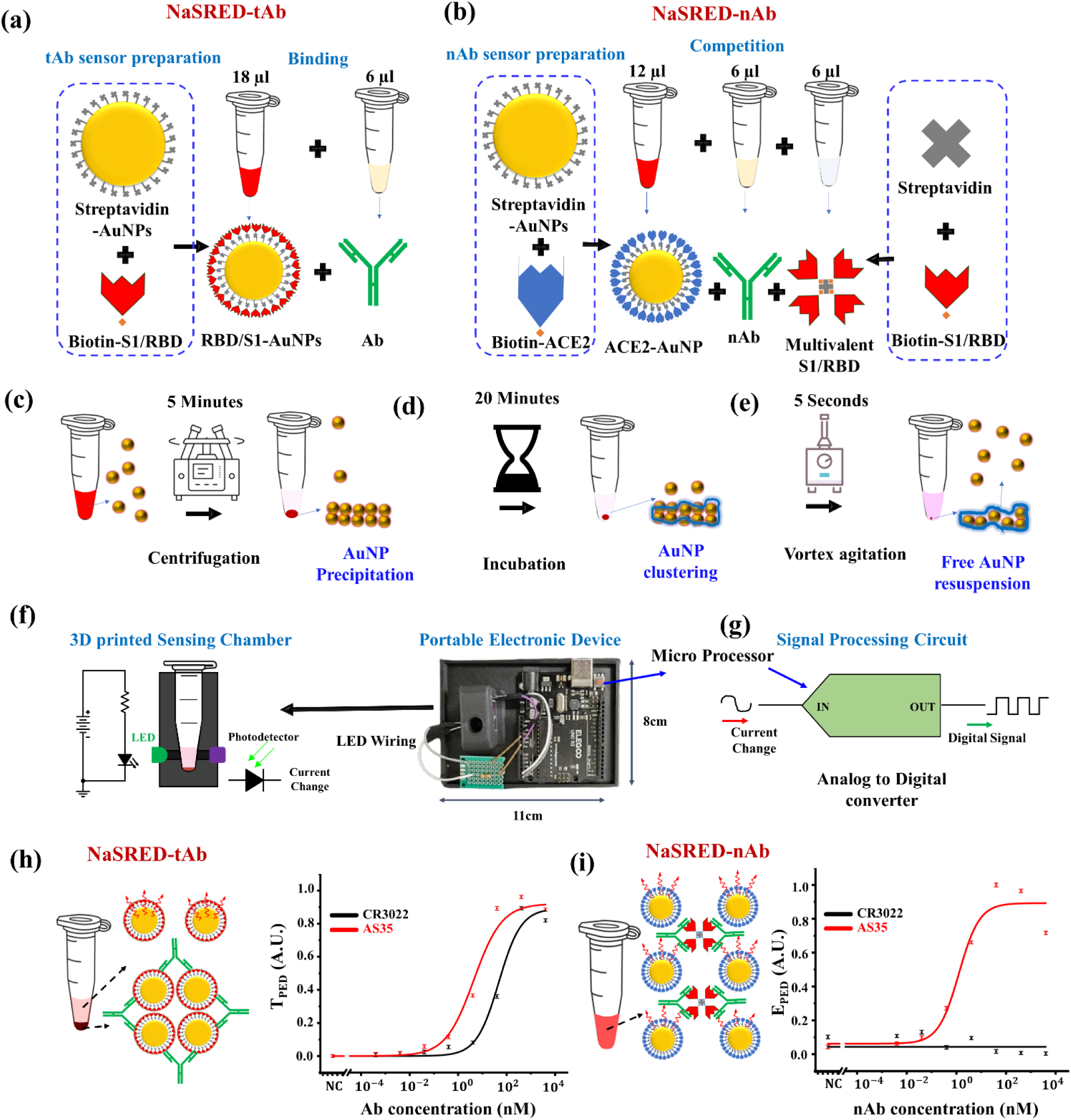
The design scheme of NasRED digital sensors. (a) NasRED-tAb (total antibody) sensing preparation: RBD (or S1)-AuNPs formation by streptavidin-biotin reaction, and then simple mixing with Ab-containing sample. (b) NasRED-nAb (neutralizing antibody) sensing preparation: ACE2-AuNPs formation by streptavidin-biotin reaction and mixing of ACE2-AuNPs, nAb-containing sample, and streptavidin-formed multivalent S1 (or RBD) molecules. The multivalent-S1 (or RBD) molecules are created by mixing biotinylated RBD or S1 with streptavidin and used to evaluate the competition of ACE2-AuNPs and nAbs for binding. (c-e) Rapid detection process comprising centrifugation of the mixtures to create a high-concentration region for accelerated reaction, followed by 20 minutes of incubation time allowing the stable clusters formation, and subsequently resuspension of the non-reacting AuNPs by vortex agitation. (f,g) Key components of the portable electronic readout system comprise an LED, a photodetector, a tube chamber, and (g) signal digitalization customized circuitry. (h-i) Representative data highlight the differences between NaSRED-tAb and NasRED-nAb assays: (left) schematics of Ab-modulated AuNP cluster distributions, where Abs trigger the formation of AuNP clusters for tAb assay but nAbs block streptavidin-RBD proteins from binding to ACE2-AuNPs; (right) sensing curves plotting normalized optical signals (transmission *T*_*PED*_ for tAb and extinction *E*_*PED*_ for nAb) against AS35 (red) and CR3022 (black) using WT-RBD-functionalized AuNP sensors for tAb and ACE2 functionalized AuNP sensors and multivalent WT-RBD for nAb. NC stands for negative control. *T*_*PED*_ and *E*_*PED*_ were obtained by averaging signals collected along five different orientations in the PED tube holder for each sample (Methods section).

Although they utilize the same PED system, the total Ab assay (NasRED-tAb) and the neutralization assay (NasRED-nAb) differ in AuNP sensor design and functionality. While NasRED-tAb is based on the Ab binding to S protein (or RBD) (**Fig. 1a**), similar to our demonstrated antigen and antibody sensing methods^19–22^, NasRED-nAb analyzes the ability of nAbs to compete with ACE2 binding to the S protein (or RBD) (**Fig. 1b**). In NasRED-tAb, AuNPs are coated with the S protein (or RBD) through streptavidin-biotin binding to act as S-AuNP or RBD-AuNP tAb sensors (**Fig. S2**). Upon mixing with commercially available, high-affinity RBD-binding human antibodies (*e.g*., AS35 (from Acrobiosystems) and CR3022 (from Absolute antibody), the AuNP-based tAb sensors bind to the antibodies and bridge into clusters, eventually precipitating to the bottom of the microcentrifuge tube. Consequently, the amount of free-floating S (or RBD)-AuNP sensors and their plasmonic extinction signals decrease (increased optical transmission) with the antibody concentration and affinity. The limit of detection (LoD) for the AS35 and CR3022 were determined as ∼3 pM and ∼15 pM by finding the lowest concentration where the PED signal significantly differs from the background (details in Methods section), respectively, due to their comparably high binding affinity (dissociation constants (*K*_*D*_s) of 0.14 nM and 6.3 nM, respectively)^23^ to the wild-type SARS-CoV-2 S protein RBD (**Fig. 1h**).

In NasRED-nAb, in contrast, AuNPs are coated with ACE2 (or ACE2-AuNPs) through streptavidin-biotin binding. Uniquely, a multivalent molecule representing the antigen of the viral variant of interest is introduced as a second protein in the sensing solution. Such a multivalent antigen, for example, can be a dimerized RBD or S protein from commercial sources or, alternatively, an artificially created multimer by binding biotinylated RBD or S proteins to streptavidin. Without nAbs that can interfere with the RBD (or S) to ACE2 binding (*e.g*., at very low concentrations of AS35 or any concentration of CR3022)^24^, the AuNPs precipitate and, as a result, produce low optical extinction. However, when high-potency nAbs are present at sufficient concentration to attach to the RBD (or S) and thus prevent their binding to ACE2, the amount of free-floating ACE2-AuNPs and their optical extinction signals remain mostly unchanged. Therefore, distinct from NasRED-tAb, the extinction increases with the nAb concentration and potency of AS35 (**Fig. 1i**).

## Detection of monoclonal antibodies (mAbs) by NasRED-tAb

Antibody binding to viral antigens is a prerequisite for virus neutralization, and differentiation of the ability of Abs to bind variants is of high interest. Because of mutations in key epitopes, viral antigen variants differ in their binding affinities to Abs, which should accordingly affect the NasRED-tAb signals measured using AuNPs functionalized with different antigen variants. Here, we tested a panel of neutralizing mAbs against RBD of SARS-CoV-2 variants Wuhan-Hu-1 (wild-type or WT), Gamma (B.1.1.28.1, or 20J/501Y.V3 or P1^25^), and Omicron (BF.7 or BA.5.2.1.7 and BA.4.6^26^) (**Fig. 2**). The inhibitory activity of these mAbs, derived from immunized mice, had been described previously using a focus reduction neutralization test (FRNT) on Vero cells with the WA1-2020 isolate^27^. The mAbs all showed strong binding to the S protein or its RBD of the WT variant as measured by ELISA; however, they differed in their neutralization potency: SARS2-02, SARS2-38, and SARS2-71 have a low half-maximum effective concentration (EC50) of 4, 5, and 8 ng/mL determined by FRNT, correspondingly, whereas SARS2-03, SARS2-10, and SARS2-31 have higher EC50 values of 670, 694 and 246 ng/mL, respectively (**Table 1**)^27^. All mAbs, except SARS2-03, effectively inhibit ACE2 binding and target different epitopes of the S protein, evidenced by their varying level of inhibition in competition tests with other structurally well-characterized human antibodies (COV2-2130, COV2-2196, COV2-2676, and CR3022)^27^.

**Figure 2.**
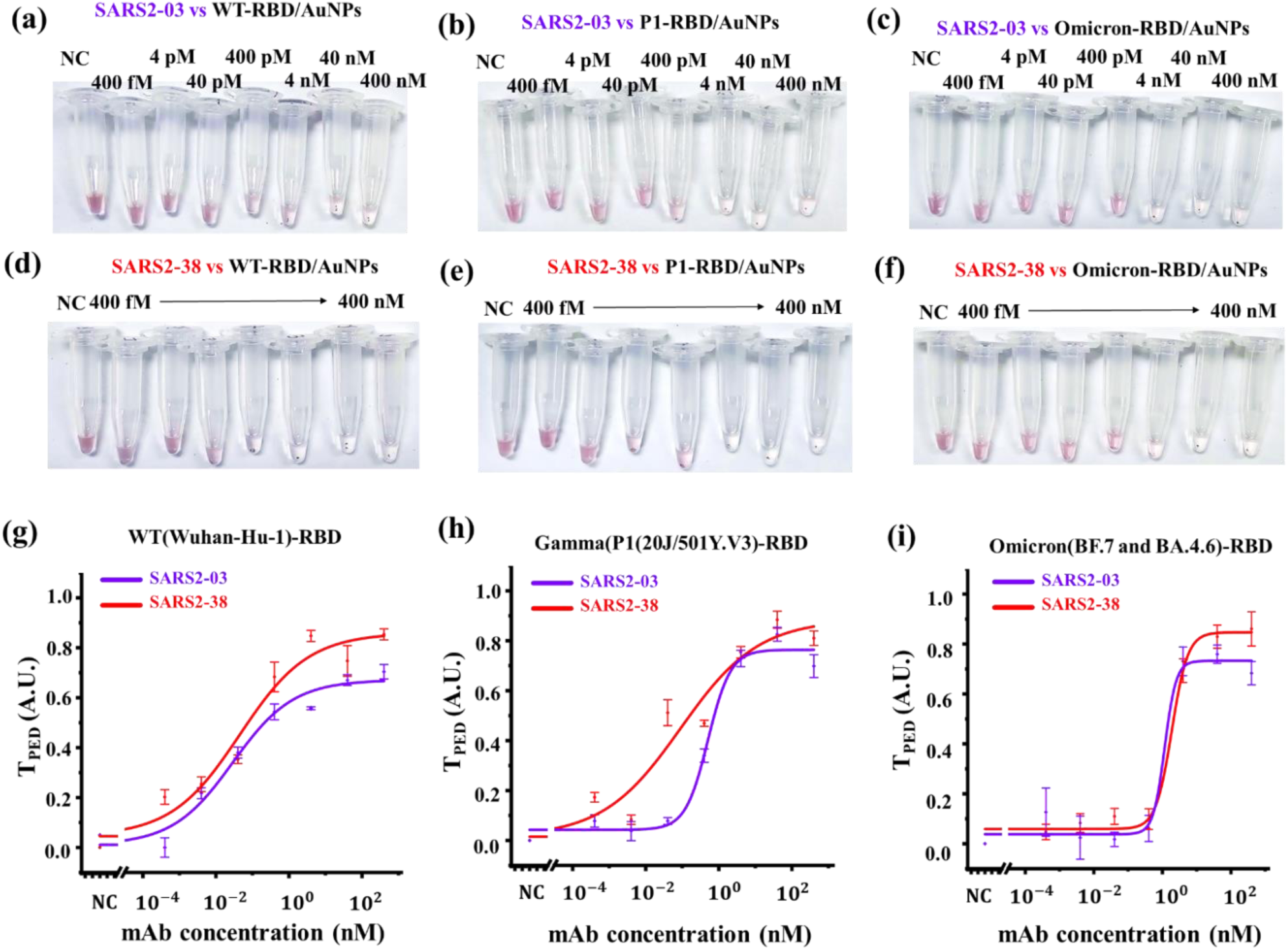
NaSRED-tAb distinguishes between the binding efficacy of monoclonal antibodies to different SARS-CoV-2 variant antigens. (a-f) Representative optical images of reaction tubes ready for electronic readout to analyze the binding of mAbs SARS2-03 and SARS2-38 to WT-RBD, P1-RBD, and Omicron-RBD functionalized AuNP sensors. The mAb concentrations are indicated on each corresponding tube. NC is Negative Control, where only PBS dilution buffer without mAb is in the testing tubes. (g-i) Sensing curves of mAbs SARS2-03 (purple lines) and SARS2-38 (red lines) against SARS-CoV-2 variant antigens (WT-RBD, P1-RBD, Omicron-RBD). Here, the optical transmission signals through the supernatant of the sensing tubes were collected on PED along five different orientations, averaged, and further normalized as *T*_*PED*_ (see Methods).

**Table 1.** Binding and inhibition efficacy of the mAbs and their EC50 values with reference Vero-TMPRSS2 FRNT data for SARS2-03, SARS2,31, and SARS2-10 based on VanBlargan et al. (2021)^27^

| Antibody | ELISA reactivity |  |  | Neutralization EC50 (ng/mL) | % ACE2 inhibition | % reference mAb binding inhibition |  |  |  |
| --- | --- | --- | --- | --- | --- | --- | --- | --- | --- |
|  | Spike | NTD | RBD |  |  | COV2-2130 | COV2-2196 | CR3022 | COV2-2676 |
| SARS2-02 | 4 | 0.1 | 4 | 4 | 100 | 100 | 100 | 14 | 3 |
| SARS2-38 | 4 | 0.1 | 4 | 5 | 99 | 95 | 1 | 12 | 3 |
| SARS2-71 | 4 | 0.1 | 4 | 8 | 100 | 2 | 99 | 8 | 5 |
| SARS2-03 | 4 | 0.1 | 4 | 670 | 4 | 5 | -2 | 68 | 4 |
| SARS2-10 | 4 | 0.1 | 4 | 694 | 100 | 1 | 5 | 97 | 12 |
| SARS2-31 | 4 | 0.1 | 4 | 246 | 100 | 1 | -1 | 98 | 0 |

The optical images of testing tubes for mAbs SARS2-03 and SARS2-38 (**Fig. 2a-f**, additional images available in **Fig. S3**) generally showed more transparent solutions at higher mAb concentrations, which is attributed to increased AuNP sensor precipitation. In comparison, the pelleted complexes displayed visual color differences for the different SARS-CoV-2 RBD variants, particularly at moderate to low mAb concentrations (*i.e*., pM to nM range), and therefore could be used to qualitatively visualize the ability of the mAbs binding to different variant antigens. For signal quantification, the tubes were analyzed by PED to attain sensing curves and calculate their corresponding LoDs^21^. The signals for the tAb tests, *T*_*PED*_, were normalized to a range between 0 and 1 for semiquantitative and comparative binding efficacy analysis of different mAbs to variant viral antigens, using a clear buffer solution as a positive control (PC) and a mixture of AuNP sensing solution with buffer (but without mAbs) as a negative control (NC) (^*T*^*PED* approaches 1 at 100% AuNP precipitation, Method section). These experiments showed that mAbs SARS2-38 and SARS2-03 produced high ^*T*^*PED* signals against all viral variants that were tested (*e.g*., ∼0.8 or higher for SARS2-38 and ∼0.6 or higher for SARS2-03), indicating a high mAb binding efficacy and, accordingly, a very high level of AuNP precipitation. These findings were consistent with previous analyses using ELISA^27^. Comparatively, the slightly lower ^*T*^*PED* values for SARS2-03 and higher NasRED-tAb EC50 values in its corresponding dose-response curves (Table 2) compared to SARS2-38 suggested that SARS2-03 had lower binding efficiency to the tested RBD proteins.

**Table 2.** NasRED-tAb analysis of mAbs against different SARS-CoV-2 variants

| Model Parameters | Antigen | SARS2-38 | SARS2-03 |
| --- | --- | --- | --- |
| $T_{\max}$ | WT-RBD | 0.85 | 0.67 |
|  | Omicron-RBD | 0.84 | 0.73 |
|  | P1-RBD | 0.88 | 0.76 |
| Slope | WT-RBD | 0.46 | 0.52 |
|  | Omicron-RBD | 1.86 | 2.6 |
|  | P1-RBD | 0.4 | 1.4 |
| EC50 (ng/mL) | WT-RBD | 6.85 | 4.42 |
|  | Omicron-RBD | 279 | 172 |
|  | P1-RBD | 13.6 | 77 |
| LoD (pM) | WT-RBD | $4 \times 10^{-3}$ | $4.2 \times 10^{-1}$ |
| | Omicron-RBD | 498 | $1.2 \times 10^3$ |
| | P1-RBD | $5.7 \times 10^{-2}$ | 134 |

Overall, the NasRED-tAb results (**Fig. 2g-i**, **Table 2**) demonstrated a femtomolar sensitivity to quantify the two mAbs over a broad (∼ 6 log10) detection range, with the LoD values of the mAbs determined from the ^*T*^*PED* plots as 4 fM for SARS2-38 against WT-RBD but 1.2 nM for SARS2-03 against Omicron-RBD (**Fig. 2g, i**). As an in-solution immunological assay, the prozone phenomenon (or hook effect at high dose)^28^ was also observed in some cases in the NasRED-tAb experiments (**Fig. 2 g-i**), where the NasRED signal *T*_*PED*_decreased slightly at the highest mAb concentrations (*e.g*., ∼100 nM or above). This was attributed to a decrease in AuNP cluster formation, resulting from higher surface coverage of the RBD-functionalized AuNPs by mAbs, which competed against mAb-induced inter-particle binding responsible for AuNP cluster formation. To minimize this impact, a rational design of the antigen density on the AuNPs and more precise calibration of the reagent concentrations could be beneficial.

## NasRED-nAb sensor design

Although measuring the mAb binding to a variant viral antigen is valuable, it does not provide insight about their capability to neutralize S or RBD receptor binding. This is exemplified by the lack of neutralization by CR3022 (**Fig. 1**) and SARS2-03 (**Fig. 2**, **Table 1**) despite their avid binding to SARS-CoV-2 RBD proteins. Because one crucial mechanism of mAb-mediated neutralization of SARS-CoV-2 is through inhibition of the viral S protein binding to the human ACE2 receptor, we conceived the idea of a three-component competitive assay for neutralization analysis, NasRED-nAb^29^, comprising ACE2-functionalized AuNPs as signal reporters, streptavidin-conjugated multivalent S (or RBD), and nAbs to be analyzed. Unlike the NasRED-tAb design, the NasRED-nAb signals depend on the competition between the nAbs and the S (or RBD) protein binding to the ACE2 receptor for any virus variant.

To explore the impact of reagent concentration on the nAb sensing performance for assay optimization, we first studied the reactivity between the multivalent WT-RBD (fixed at 500 pM) and ACE2-AuNPs of different concentrations from 0.05 to 0.8 *OD*_*PED*_ (**Fig. 3a**). Here *OD*_*PED*_ was defined as the PED-measured optical density, representing the ACE2-AuNP extinction normalized in reference to a buffer solution (Methods). Following the same NasRED-tAb processing steps (centrifugation, incubation, and vortex agitation, **Fig. 1c-e**) described above, samples were measured using the PED device to determine the optical signal difference (**Fig. 3c**, blue curve). The results showed that ACE2-AuNps at *OD*_*PED*_∼0.4 produced the largest color contrast for experimental visualization. To optimize the stoichiometric ratio of ACE2-AuNPs and multivalent-RBD molecules, we chose two AuNP concentrations at *OD*_*PED*_ ∼0.8 (**Fig. 3d-f**) and *OD*_*PED*_ ∼0.4 (**Fig. 3g-i**) to measure multivalent-RBD (0 to 17 nM, and 0 to 5 nM, respectively) binding. The tube colors (**Fig. 3d**) were visually almost transparent at ∼2 nM or higher multivalent-RBD for ACE2-AuNP sensors at *OD*_*PED*_∼0.8, indicating a high degree of ACE2-AuNPs precipitation. Similarly, a reaction with ∼1 nM multivalent-RBD resulted in high transparency for ACE2-AuNPs at *OD*_*PED*_ ∼0.4 (**Fig. 3g**). This RBD-concentration-dependent signal transition could be used as a reference to optimize the concentrations and molar ratios of ACE2-AuNPs and RBD molecules in the nAb assay. A sufficiently high ACE2-AuNP concentration, desired to reveal effective nAb neutralization with a clear red solution color, was useful for high signal-to-noise ratio PED detection. Accordingly, the amount of RBD molecules in the nAb assay needs to be rationally designed, i.e., high enough to completely precipitate the AuNPs in the absence of nAbs to maximize the assay dynamic range but not excessive to avoid unnecessarily consumption of nAbs in free-solution reactions and subsequently lowering the assay sensitivity. However, higher concentrations of ACE2-AuNPs and RBD molecules also demand more nAbs for competitive binding to produce detectable signals, thus potentially limiting the assay sensitivity and dynamic range. Here, we found that the RBD to ACE2 molar ratio from the observed color transitioning was 1:8 in both cases to trigger significant AuNP cluster formation (see Methods).

**Figure 3.**
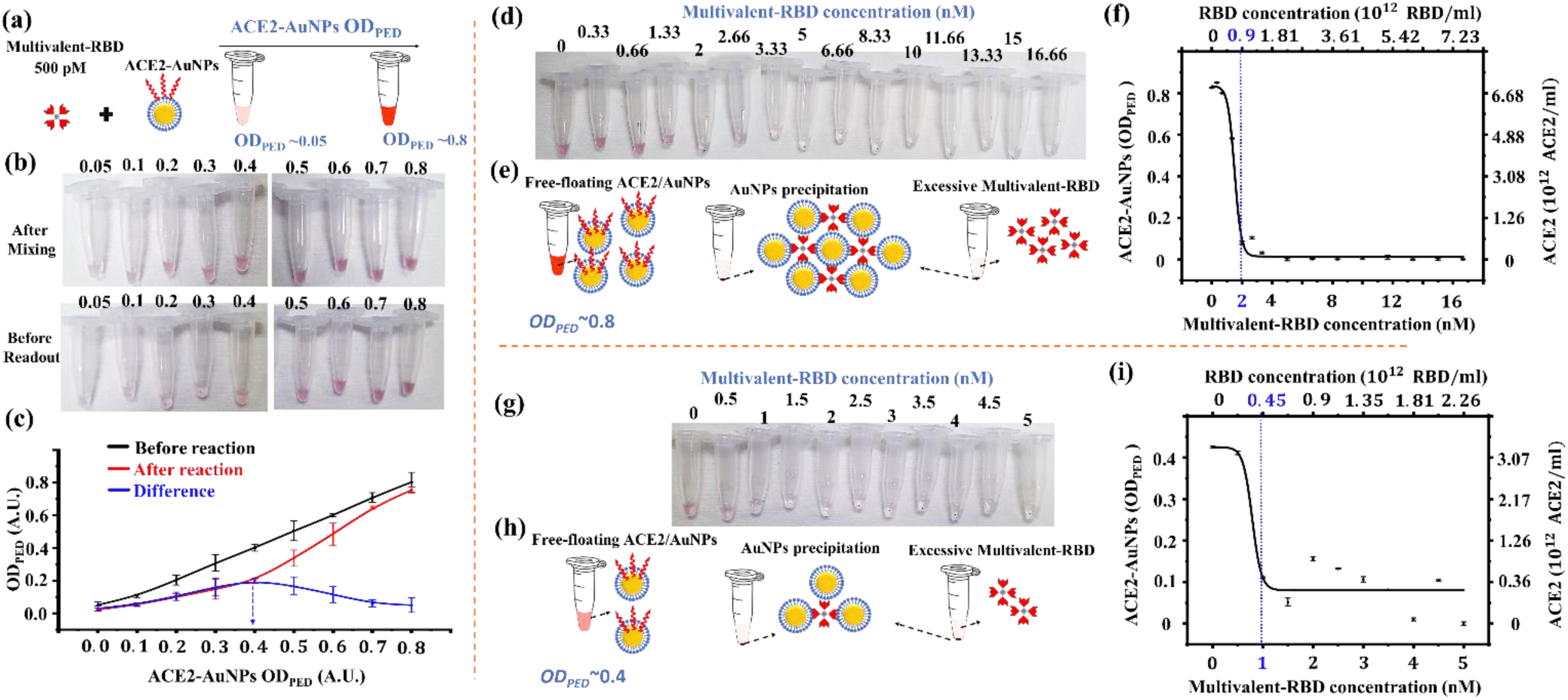
Design of reagent concentrations for NasRED-nAb sensor optimization. (a-c) The impact of the initial ACE2-AuNP concentration (optical extinction signal measured by PED readout, i.e.*OD*_*PED*_, from 0.05 to 0.8) on the signal modulation after reaction with tetrameric RBD (fixed at 500 pM): (a) the schematics showing the reacting reagents, and (b) optical images of the testing tubes after mixing and before readout, and (c) the measured changes in the *OD*_*PED*_ (blue) plotted with the initial (black) and final (red) *OD*_*PED*_ values of the ACE2-AuNPs, suggesting a maximum signal change around *OD*_*PED*_∼0.4. (d-i) Further quantification of the RBD concentration impact on the sensor solution color at different ACE2-AuNP initial concentrations: (d-f) *OD*_*PED*_∼0.8, and (g-j) *OD*_*PED*_∼0.4. (d,g) The optical images of microcentrifuge tubes containing tested RBD molecules after the reaction. (e,h) Schematic illustration of the reagent reaction after the reaction when the multivalent-RBD molecules were absent (left), just enough to fully precipitate the ACE2-AuNPs (middle) and excessive (right). (f, i) The sensor *OD*_*PED*_ signal plotted as a function of RBD concentration, with the corresponding ACE2 concentration (derived from the measured *OD*_*PED*_, Methods) included as the y-axis on the right. The blue dotted lines and number indicate the multivalent RBD concentrations selected for subsequent testing conditions 1 and 2. Here, the optical transmission signals through the AuNP solution tubes were collected on PED along five different orientations, averaged, and further normalized as *OD*_*PED*_ (see Methods).

From these analyses, we designed two assay conditions: condition 1 (ACE2-AuNPs at *OD*_*PED*_ ∼0.8, 2 nM multivalent-RBD) and condition 2 (ACE2-AuNPs at *OD*_*PED*_∼0.4, 1 nM multivalent-RBD), for the NasRED-nAb assay (**Fig. 4a,b**). Using commercial nAb (AS35) as an example against WT-RBD under both conditions, three replicate experiments were performed using the same test protocol (5 minutes of 1,200× g centrifugation, 20 min of incubation, and 5 sec of vortex agitation at 2050 RPM) as described above. The nAb sensing signal *E*_*PED*_, calculated from the transmission signal of the Ab sample and normalized against buffer and AuNP sensing solutions (Methods), were reproducible (**Fig. 4c-e**, **Table 3**), but were different for the two test conditions. The neutralization concentration of AS35 to modulate the signal *E*_*PED*_ by 50%, i.e., the EC50 on NasRED-nAb assay (or NasRED-nAb EC50) was >3 fold lower in condition 2 (1,450 ng/mL) as compared to condition 1 (4,680 ng/mL) and more comparable to the manufacturer-provided hACE2-binding ELISA EC50 (1,320 ng/mL, Acrobiosystems) (**Fig. 4c-e**).

**Figure 4.**
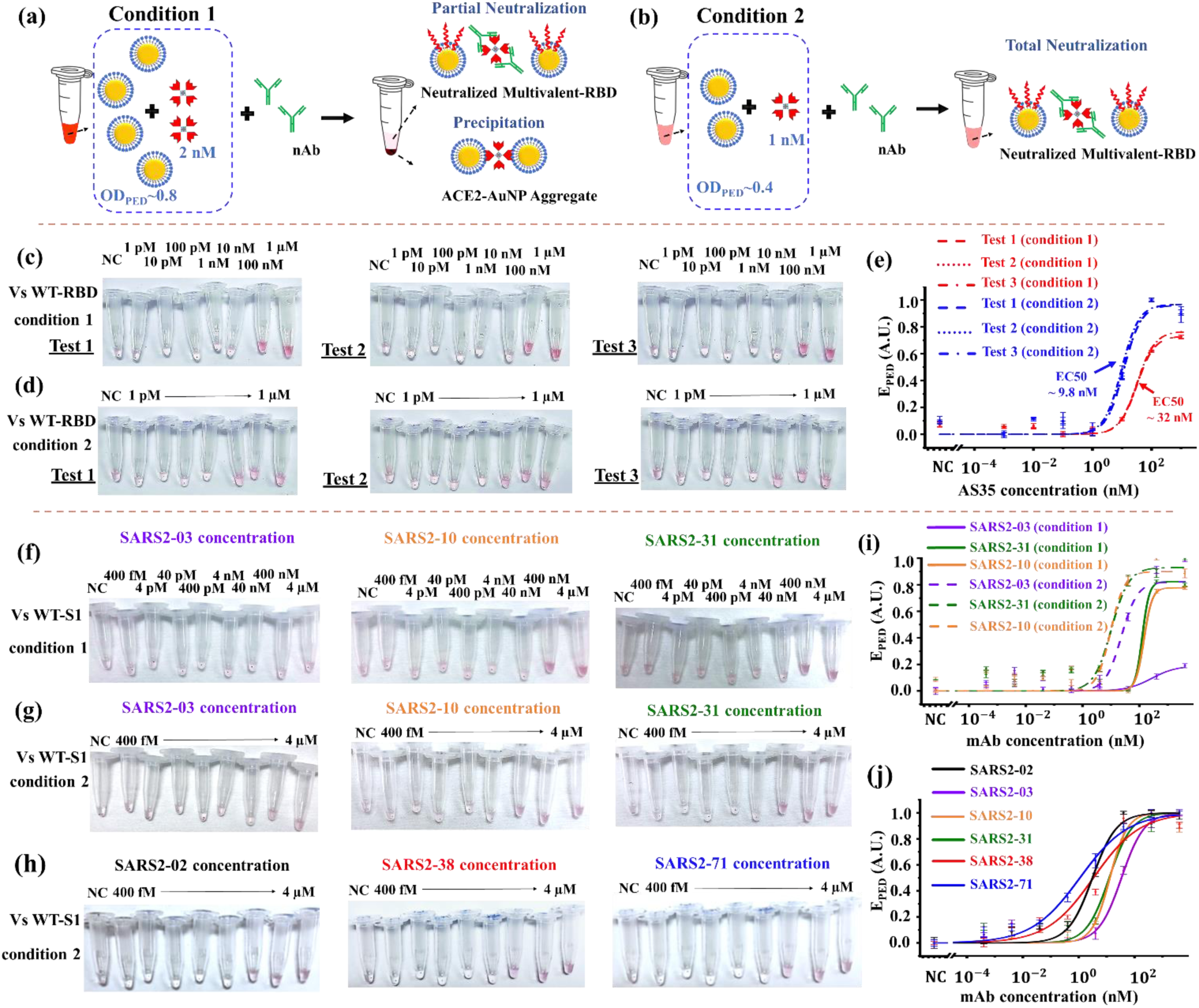
NasRED-nAb distinguishes mAbs neutralizing potencies against viral antigens under different testing conditions. (a,b) Schematic representation showing the two testing conditions affect the available free-floating AuNPs and the NaSRED-nAb signal. (c,d) Optical images of NASRED-nAb test tubes to evaluate mAb AS35 against WT-RBD in PBS under both conditions 1 and 2 in three replicate tests, where three sets of samples were prepared and tested using the same protocol, to evaluate the nAb test reproducibility. (e) Normalized optical extinction signals *E*_*PED*_ under conditions 1 (red lines) and 2 (blue lines), with the EC50 values marked. (f-h) Optical images of the testing tubes for different mAbs against WT-S1 in PBS: (f) SARS2-03, SARS2-10, and SARS2-31 tested under condition 1, (g) SARS2-03, SARS2-10, and SARS2-31 tested under condition 2, and (h) SARS2-02, SARS2-38, and SARS2-71 tested under condition 2. (i-j) Normalized optical extinction *E*_*PED*_ plotted versus the mAb concentrations: (i) SARS2-03, SARS2-10, and SARS2-31 under condition 1 (solid lines) and condition 2 (dashed lines), and (j) all the mAbs under condition 2. Buffer without mAbs was used as Negative Control (NC). Here, the optical extinction signals through the supernatant of the nAb sensing tubes were collected on PED along five different orientations, averaged, and further normalized as *E*_*PED*_ (see Methods).

**Table 3.** Fitting parameters between repeated nAb assays. Here, the slope, Emax, and most importantly, the EC50 values are closely consistent, showing the neutralizing assay’s reproducibility in quantitative measurements of a triplicate for AS35 neutralization against WTRBD.

| Model Parameters | Test condition 1 |  |  | Test condition 2 |  |  |
| --- | --- | --- | --- | --- | --- | --- |
|  | Test 1 | Test 2 | Test 3 | Test 1 | Test 2 | Test 3 |
| $E_{\max}$ | 0.768 | 0.731 | 0.731 | 0.968 | 0.961 | 0.969 |
| EC50 ($10^3$ ng/mL) | 4.89 | 4.75 | 4.72 | 1.45 | 1.69 | 1.74 |
| Slope | 1.43 | 1.48 | 1.47 | 1.48 | 1.47 | 1.47 |
| R-Square (COD) | 0.99991 | 0.99992 | 0.99991 | 0.99973 | 0.99975 | 0.99973 |

Next, low-potency neutralizing mAbs (SARS2-03, SARS2-10, and SARS2-31) were tested against the S1 subunit of the WT-spike protein under conditions 1 and 2 (**Fig. 4f,g**), and high-potency mAbs (SARS2-02, SARS2-38, and SARS2-71) were tested under condition 2 (**Fig. 4h**). The WT-S1 subunit allows for evaluation of mAb binding both to and outside the RBD. For example, SARS2-38, like mAb CoV2-2130^27^, directly overlaps the RBD binding site for ACE2, thus inhibiting RBD binding to ACE2. In contrast, SARS2-03 overlaps with mAb CR3022, which engages the base of the RBD site distal from the receptor binding site and does not inhibit RBD to ACE2 binding (**Table 1**)^27^. It is known that CR3022 recognizes an epitope that is only accessible when the RBD is in a specific conformation, and its constant region interferes with the N-terminal domain of the S protein^30^. Analyzing the impact of testing conditions (**Fig. 4i**), the observed NasRED-nAb EC50 values were found to be 10 to 20 times higher under condition 2 compared to condition 1 (**Fig. 4i**, **Table 4**). The approximately two-fold difference in reagent (ACE2-AuNP and RBD) concentrations caused a much more pronounced impact on the NasRED-nAb EC50 values for the low-potency mAbs (**Fig. 4i**) than high-potency mAb AS35 (**Fig. 4e**). This observation can be understood from the perspective of the competitive protein binding mechanism of our assay, where the competition between multivalent proteins^31,32^, including streptavidin-bound multivalent RBD, AuNP-carried ACE2, and bivalent nAbs, depend heavily on the protein concentration, different binding epitopes, and binding affinities. For example, the binding affinity between RBD and ACE2 (*K*_*D*_ 15.2 nM)^33,34^ is comparable to that between the RBD and AS35 (*K*_*D*_ 5.1 nM)^35^ or SARS2-38 (*K*_*D*_ 6.5 nM)^27^. Therefore, the reagent concentration effect was less obvious for such high-potency mAbs since their higher binding affinity played the primary role in outcompeting RBD for binding to ACE2^35^. These studies suggested the importance of NasRED-nAb assay design parameters in detecting samples with significant heterogeneity in neutralization potencies. Furthermore, the mAb neutralization potency was quantified by NasRED-nAb using the same WT-S1 protein, and the low-potency mAbs (SARS2-03, SARS2-10, and SARS2-31) had EC50 values (1,000 to 7,000 ng/mL) about one order of magnitude higher than of the high-potency mAbs (SARS2-02, SARS2-38, and SARS2-71, EC50 100 to 500 ng/mL, **Table 4**). This observation was consistent with our prior FRNT virus neutralization studies (**Table 1**)^27^, suggesting effective NasRED-nAb quantification.

**Table 4.** Design impact on NasRED-nAb performance in analyzing neutralization capabilities of mAbs against WT-S1 as the SARS-CoV-2 variant. Condition 1: ACE2-AuNP at ^*OD*^_*PED*_∼0.8, multivalent S1 at ∼2 nM. Condition 2: ^*OD*^_*PED*_∼0.4, multivalent S1 at ∼1 nM.

| Model Parameter | Test condition | SARS2-02 | SARS2-03 | SARS2-10 | SARS2-31 | SARS2-38 | SARS2-71 |
| --- | --- | --- | --- | --- | --- | --- | --- |
| $E_{max}$ | Condition 1 | -- | 0.19 | 0.78 | 0.82 | -- | -- |
|  | Condition 2 | 1 | 0.82 | 0.9 | 0.93 | 1 | 1 |
| Slope | Condition 1 | -- | 1.00 | 3.2 | 3.5 | -- | -- |
|  | Condition 2 | 1.01 | 1.54 | 1.75 | 1.41 | 0.52 | 0.50 |
| EC50<br>($10^3$ ng/mL) | Condition 1 | -- | 80.4 | 21.6 | 18.88 | -- | -- |
|  | Condition 2 | 0.46 | 3.24 | 1.21 | 1.52 | 0.5 | 0.16 |

## NasRED-nAb differentiating mAbs potency against viral variants

To further validate NasRED-nAb in analyzing the neutralizing potency of mAbs, four mAbs (SARS2-02, SARS2-03, SARS2-38, and SARS2-71) were tested against different SARS-CoV-2 RBD antigens, including WT-RBD, P1-RBD, and Omicron (BF.7 and BA.4.6)-RBD (**Fig. 5**). As expected, the three high-potency mAbs (SARS2-02, SARS2-38, and SARS2-71) were neutralizing against WT-RBD under the test condition 1 (**Fig. 5a**, **Table 5**) with EC50 values from 2,000 to 10,000 ng/mL. Although the NasRED-EC50 values were about 3 orders of magnitude higher than that determined by FRNT (4 to 8 ng/mL) as a result of the high reagent concentration effect discussed above, NasRED and FRNT data were consistent in comparing the relative neutralizing activity of the different mAbs, suggesting SARS2-02 was slightly more neutralizing than the others and SARS2-71 was the least potent with the largest EC50 value. Furthermore, these three high-neutralizing mAbs were less effective against P1-RBD, with the NasRED-EC50 concentrations about one order of magnitude higher than measured of WT-RBD (**Fig. 5b**, **Table 5**). This diminished neutralization was expected considering key mutations on the P1-RBD, including K417T, E484K, and N501Y. In particular, SARS2-02 was very sensitive to the E484K^27^ mutation, whereas SARS2-38 and SARS2-71 were not, and the NasRED-nAb indeed demonstrated that SARS2-02 almost completely lost neutralizing activity, whereas SARS2-38 and SARS2-71 maintained their neutralizing capability. In comparison to these three high-potency mAbs, SARS2-03 did not produce appreciable neutralization within the tested concentration range against either RBD variant because this mAb does not block ACE2 to RBD binding^27^.

**Figure 5.**
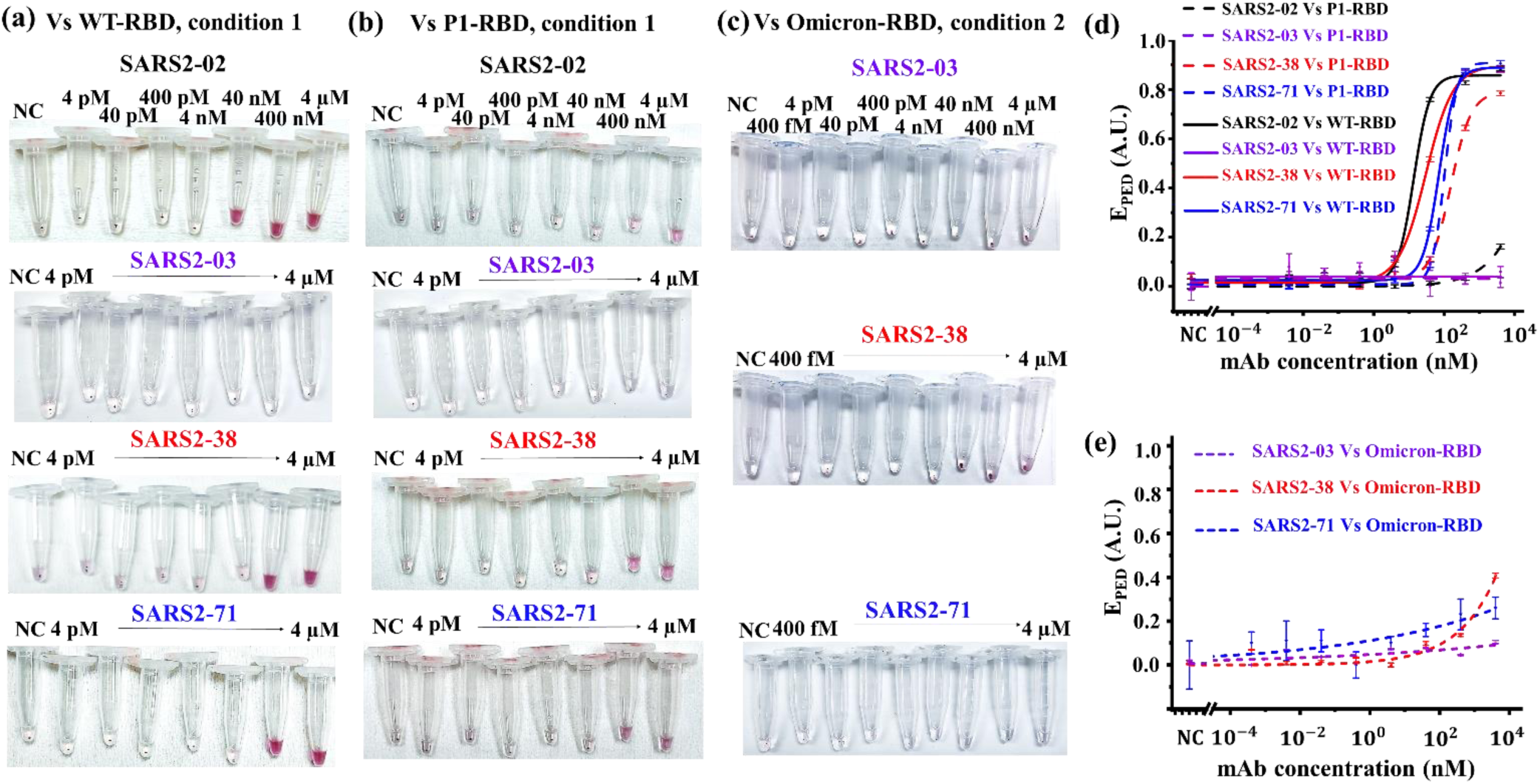
NaSRED-nAb quantitatively discriminates mAbs’ potency against different variants. (a,b) Optical images of testing tubes and (d) normalized nAb signal *E*_*PED*_ of tested mAbs SARS2-02, SARS2-03, SARS2-38, and SARS2-71 against WT-RBD (solid lines) and P1-RBD (dash lines) under condition 1, indicating reduced neutralizing potency against the P1 variant. (c, e) Optical images of testing tubes and (e) normalized nAb signal *E*_*PED*_ of tested mAbs SARS2-03, SARS2-38, and SARS2-71 against omicron RBD under condition 2, indicating low neutralizing potency of such mAbs against the Omicron variant. Here, the optical extinction signals through the supernatant of the sensing tubes were collected on PED along five different orientations, averaged, and further normalized as *E*_*PED*_ (see Methods).

**Table 5.** Quantitative differentiation between mAbs’ potency across WT and P1 variants under Condition 1: ACE2-AuNP at ^*OD*^_*PED*_∼0.8, multivalent S1 at ∼2 nM.

| Model Parameter | Antigen | SARS2-02 | SARS2-38 | SARS2-71 | SARS2-03 |
| --- | --- | --- | --- | --- | --- |
| $E_{max}$ | WT-RBD | 0.89 | 0.88 | 0.88 | NN* |
|  | P1-RBD | 0.09 | 0.78 | 0.91 | NN |
| Slope | WT-RBD | 1.7 | 1.11 | 1.88 | NN |
|  | P1-RBD | 0.02 | 1.42 | 2.43 | NN |
| EC50<br>(ng/mL) | WT-RBD | $2.13 \times 10^3$ | $4.08 \times 10^3$ | $1 \times 10^4$ | NN |
| | P1-RBD | $1.8 \times 10^7$ | $2.07 \times 10^4$ | $1.5 \times 10^4$ | NN |

\*NN: SARS2-03 showed no neutralization against WT-RBD and P1-RBD in NasRED-nAb condition 1 design.

**Table 6.** mAbs’ neutralizing potency against Omicron-RBD under Condition 2: ^*OD*^_*PED*_∼0.4, multivalent S1 at ∼1 nM.

| Model Parameter | SARS2-38 | SARS2-71 | SARS2-03 |
| --- | --- | --- | --- |
| $E_{max}$ | 0.41 | 0.26 | 0.1 |
| Slope | 1.05 | 0.1 | 0.16 |
| EC50 (ng/mL) | $8.05 \times 10^4$ | $6.78 \times 10^8$ | $2.3 \times 10^{12}$ |

Further, we compared SARS2-38 and SARS2-71, two potent mAbs that were more resistant to RBD mutations, with SARS2-03, in neutralization against Omicron RBD (BF.7 and BA.4.6)^10,26,33^, which contains a larger number of mutations (*e.g*., G339D, R346T, S371F, S373P, S375F, T376A, D405N, R408S, K417N, N440K, L452R, S477N, T478K, E484A, F486V, Q498R, N501Y, and Y505H). Considering the expected loss in mAb neutralization capabilities, mAbs were tested against Omicron RBD under condition 2, i.e., at a reduced antigen density compared to condition 1, to better analyze the mAb effect within the achievable concentration ranges. We observed no neutralization signal except at the highest concentrations (4 *μ*M) of SARS2-38 and SARS2-71, indicating the ACE2-AuNPs were still tightly bound to the multivalent-RBDs despite high mAb concentrations (**Fig. 5c,e)**. This was not surprising because SARS2-71 loses neutralization capability against mutations at S477, T478, and F486^27^. In comparison, while SARS2-38 was found resistant to mutations such as L452R and E484K, its EC50 values against B.1.1.529 Omicron variant S protein increased >1,000 times in both Vero-TMPRSS2 and Vero-hACE2-TMPRSS2 cells^10^. This is likely because SARS2-38 binding to RBD is greatly diminished by mutation at position N440K^10^. In addition, the Omicron RBD has a much higher binding affinity compared to WT-RBD for the ACE2 receptor, e.g., *K*_*D*_∼2.5 nM for Omicron BA.2 and ∼15.4 nM for G614 (WT)^33^. Therefore, it becomes significantly more challenging for mAbs to protect ACE2 from binding to Omicron variants.

Previously in the NasRED-tAb tests (**Fig. 2**), SARS2-03 and SARS2-38 showed high reactivity in binding against WT-, P1- and Omicron-RBD, with the EC50 values against WT- and P1-RBD on the order of a few ng/mL. However, strong mAb binding to the RBD does not always result in neutralization, evidenced by the fact that SARS2-03 was ineffective against all RBD variants from the NasRED-nAb tests. Clearly, NasRED-nAb is better suited to analyze the neutralization potency of mAbs. Although simple *in vitro* assays such as NasRED-nAb would not replace virus-based neutralization assays to quantify the inhibitory capabilities because many factors (*e.g*., heterogeneity in human ACE2 production or humoral response to infection or vaccines). Nonetheless, NasRED-nAb can provide insight by rapidly comparing the neutralizing activity of different mAbs in virus neutralization.

## NasRED-nAb correlates with ELISA in the neutralizing analysis of human serum samples

We performed both NasRED-nAb and hACE2 binding competitive ELISA, utilizing SARS2-71 spiked in Human Pooled Serum (HPS) samples under condition 2 (**Fig. 6a,b**) to assess the assay performance under serum matrix in comparison to competitive ELISA. The NasRED-nAb EC50 (17.3 ng/mL) was comparable to that of the ELISA (15.8 ng/mL), confirming the NasRED-nAb’s feasibility for quantitative neutralizing activity analysis. Next, we used NasRED-nAb to measure human serum neutralizing activity. For the best sensing results and based on our previous study on NasRED operational mechanisms^21^, critical parameters such as centrifugation and vortex agitation speed were slightly increased (centrifugation from 1,200× *g* to 1,600× *g* and vortex from 2,050 RPM to 2,250 RPM) to consider the higher viscosity of serum compared to PBS. We performed NasRED-nAb tests on a total of 40 (10 negatives and 30 positives) ELISA-validated human sera samples (**Fig. 6c**) from vaccinated and convalescent individuals after COVID-19 infection and healthy participants collected during Serosurvey 1 in September 2021 (**Table S1**). The NasRED-nAb EC50 data were compared to the OD450 values measured by hACE2 binding competitive ELISA and found good correlation following an exponential fitting *NasRED*_*Cal*_ = 136.26 × ln(*E*_*PED*_) + 450.27 (**Fig. 6d**), with a small residual sum of squares (*RSS* = 0.198) and a perfect R-squared (*R*^2^ = 1). This empirically derived fitting was used to calibrate the NasRED-nAb signals in reference to ELISA measurements in a linear relationship, following *NasRED*_*Cal*_ = 0.94 × *ELISA* + 11.27 (*R*^2^ = 0.93) (**Fig. 6e**). In contrast, the *NasRED*_*Cal*_ and ELISA OD450 signals completely deviated from total IgG counts based on ELISA measurements (*RSS* = 2279, *R*^2^ = 0.44), further confirming that total IgG count is not a reliable measure for neutralization, but the NasRED-nAb could be applied as a surrogate method to analyze the neutralization activity of more complex clinical sera samples.

**Figure 6.**
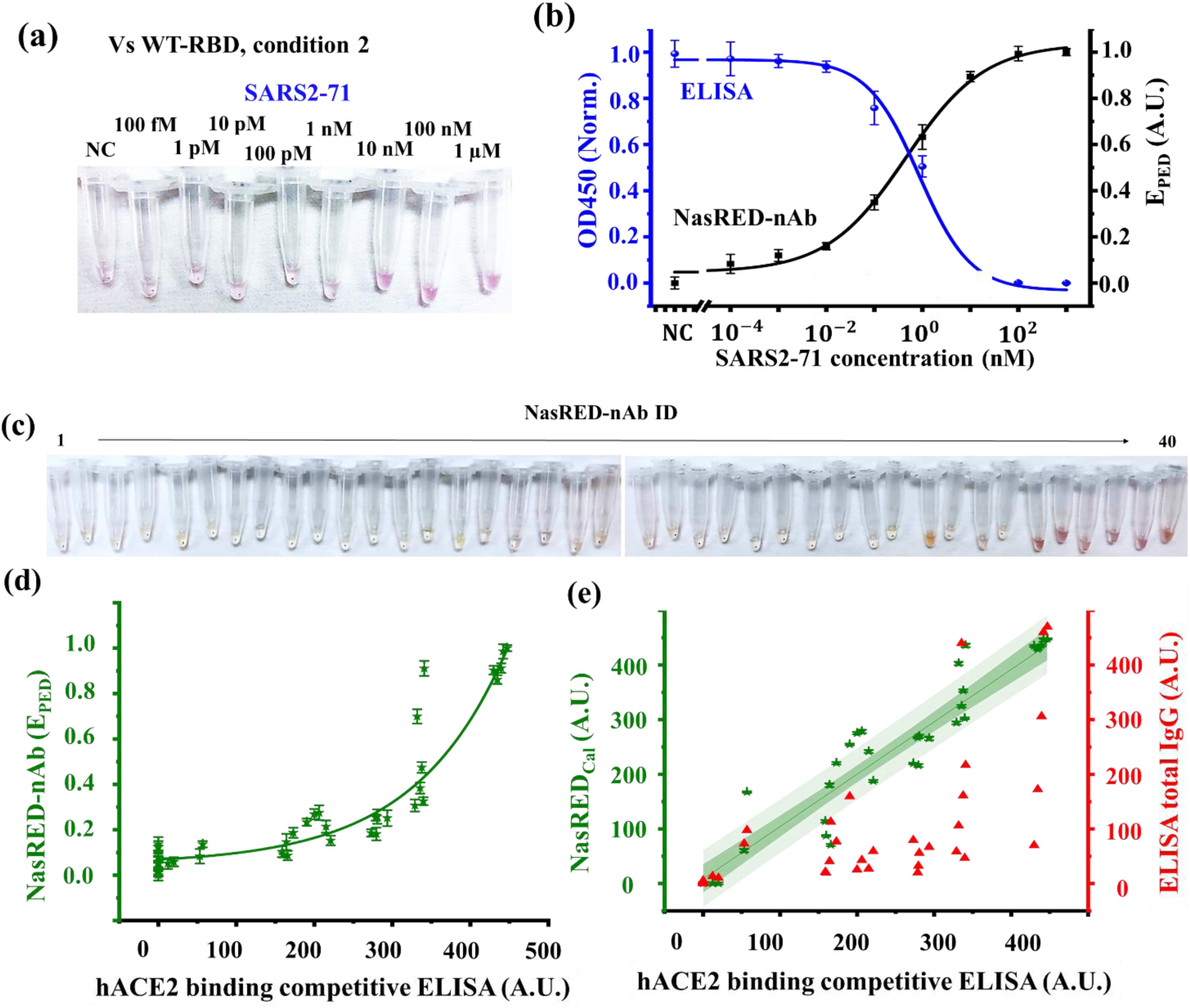
Correlation of NaSRED-nAb with hACE2 binding competitive ELISA in immunity screening. (a,b) Optical images of the testing tubes for mAb SARS2-71 spiked in HPS against WT-RBD under condition 2, and (b) the NASRED-nAb signal *E*_*PED*_ (black line) and the hACE2 binding competitive ELISA signal (normalized OD450, blue line) against SARS2-71 concentration. (c) Optical images of the testing results of 40 sera survey samples (10 negatives + 30 positives for SARS-CoV-2 wild type) tested by NasRED-nAb against WT-S1 under condition 2. (d) NasRED-nAb data (stars) and fitting curve (solid line) plotted versus the ELISA OD450 values (*R*^2^ = 1). (e) The calibrated nAb signal *NasRED*_*Cal*_ (left Y-axis, stars, and green line, with dark and light shading for the 95% confidence and prediction bands) and the total IgG counts from ELISA assay measurements (red triangles) plotted versus ELISA OD450 values, indicating a good agreement between *NasRED*_*Cal*_ and competitive ELISA (*R*^2^ = 0.93) but poor agreement with the total IgG count. Here, the optical extinction signals through the supernatant of the sensing tubes were collected on PED along five different orientations, averaged, and further normalized as *E*_*PED*_ (see Methods).

## NasRED-nAb in longitudinal seroconversion studies against WT and Omicron RBD

To analyze the total count and neutralizing potency of SARS-CoV-2-specific antibodies in serum (seroreactivity) on the NasRED platform, we utilized NasRED-tAb and NasRED-nAb to test against WT-RBD and Omicron-RBD in serum samples collected from 10 individual participants in Serosurvey 1 and 2 studies (September 2021 and October 2022) (**Fig. 7a,b**). All subjects were vaccinated against the WT spike before sample collection in Fall of 2021 and reported infection by Omicron based on symptoms before the sample collection in Fall of 2022 (**Table S2**). The signals for the tAb tests, *T*_*PED*_, were normalized as described above and in the Methods section using both WT-RBD (**Fig. 7c**) and Omicron-RBD (**Fig. 7e**). For nAb assay calibration, commercially available HPS (Innovative Research Inc.) was used as the negative control (NC), and SARS2-38 that exhibited highest neutralizing activity was spiked into HPS with the highest concentration (10 µM) as the positive control (PC). The electronic readout signal for each subject sample, *E*_*PED*_, was then normalized in reference to the NC and PC signals to a value between 0 and 1, and used to semi-quantitatively compare the neutralization capability against the WT-RBD (**Fig. 7d**) and Omicron-RBD (**Fig. 7f)**. Furthermore, the increase in tAb and nAb signals from the two surveys, Δ*T*_*PED*_ and Δ*E*_*PED*_, were also obtained from measurement with the WT and Omicron variants (**Fig. 7g,h**). The tAb and nAb signals were normalized using different standards, and therefore, their values should not be compared across the two different platforms.

**Figure 7.**
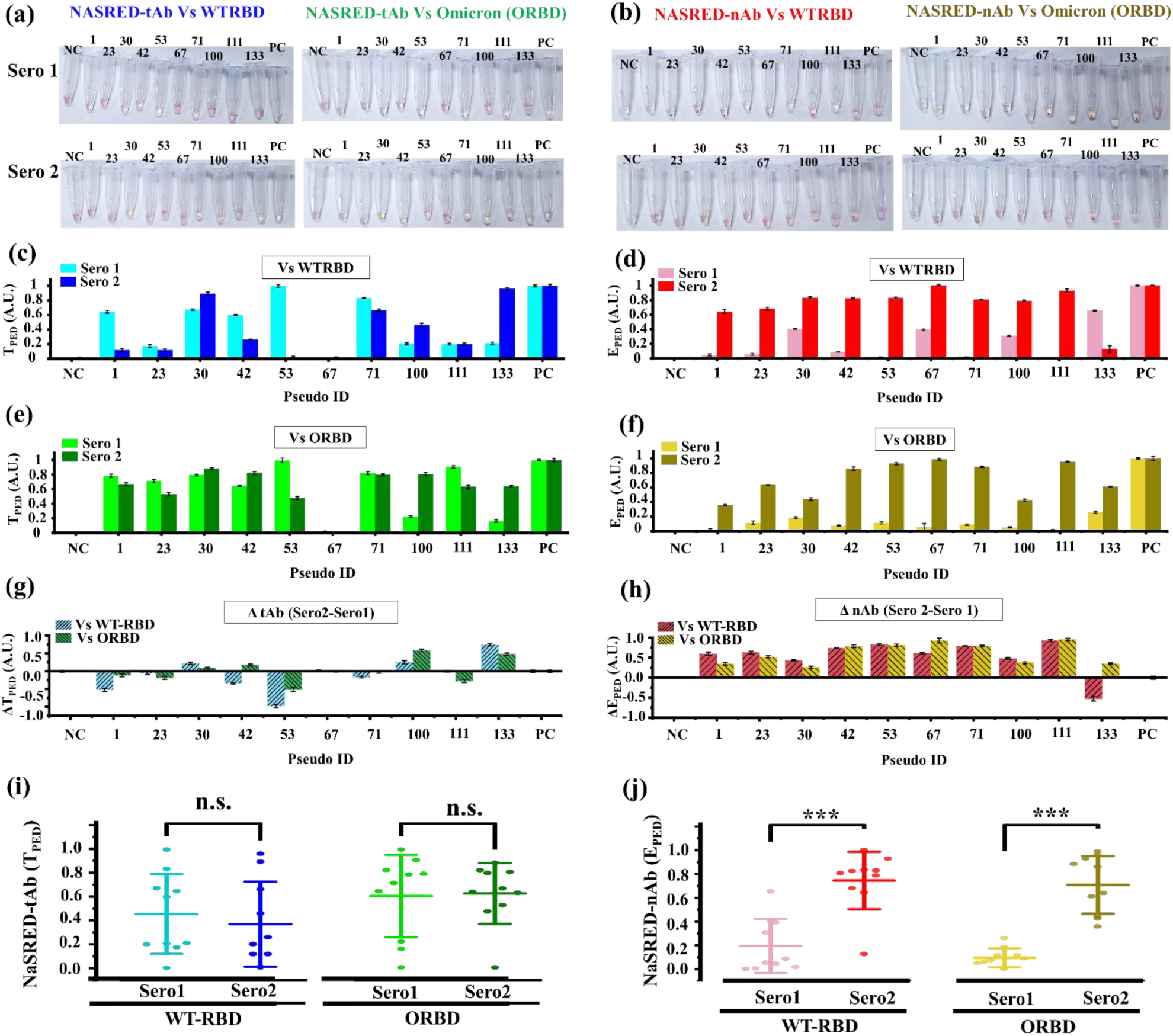
NaSRED analyzes samples in longitudinal serosurvey studies against wildtype and Omicron variants. (a,b) Optical images of the NasRED-tAb and NasRED-nAb test tubes to analyze samples collected from serosurvey 1 (post-WT vaccination 2021) and serosurvey 2 (post-omicron infection 2022) against WT-RBD/ORBD. (c,e) NaSRED-tAb and (d,f) NaSRED-nAb results plotted against the pseudoIDs of samples in serosurvey1 (light color bars) and serosurvey2 (dark color bars). (g,h) The relative changes in tAb and nAb signals between the two surveys against WT-RBD and ORBD, indicating a lack of trend in total antibody quantification but an overall increase in neutralization values. (i,j) Tukey’s two-way ANOVA analysis for the two serosurveys and variants indicated (i) a non-significant difference (P>0.05) between total antibody levels and (j) a significant difference (P<0.001) between neutralization levels. Here, the optical extinction signals through the supernatant of the sensing tubes were collected on PED along five different orientations, averaged, and further normalized as *E*_*PED*_ (see Methods).

The NasRED-tAb signal increase Δ*T*_*PED*_ was not statistically significantly for activity when measured against either the WT or Omicron variants (P > 0.05), but the nAb signal increase Δ*E*_*PED*_from the same samples did show a significant change (P < 0.001) (**Fig. 7i,j**). The observed fluctuations of the total amount of virus-binding IgG levels could be attributed to their random temporal changes, which could be affected by many factors such as allergies, inflammation, or illnesses other than COVID-19 and did not reliably predict virus neutralization. The results also indicated that the Omicron infection experienced by the subjects not only increased neutralization against Omicron variant but also greatly "back-boosted" immunity against the WT antigen by a significant amount (Δ*E*_*PED*_>0.50) (**Fig. 7d, f, h**), consistent with previous studies^36,37^. The fundamental difference between tAb and nAb tests could be clearly seen by examining representative subjects. For example, in the first survey (Fall 2021), NasRED-tAb showed that subject 53 strongly reacted (∼100%) to WT and Omicron. Still, the subject’s total IgG level was reduced significantly in the second survey (Fall 2022), particularly against the WT. In contrast, the NasRED-nAb showed that the subject had very low nAb levels against both WT and Omicron variants in survey 1 but acquired increased neutralizing capability (∼80%) in the second survey against both variants. In another example, very low levels of tAb signals were observed for subject 67 in both surveys, but appreciable increases in the nAb levels were observed against both the WT and Omicron. In summary, the study using clinical samples again proved the importance and usefulness of NasRED-nAb in neutralization studies.

## Discussion

Accurately detecting nAbs is essential for evaluating immune protection against circulating viruses, such as SARS-CoV-2, especially as new variants emerge. Traditional tAb assays, including ELISA^38^, quantify overall antibody levels but do not address whether these antibodies can neutralize virus. The NasRED-nAb platform is designed based on our recent demonstration of NasRED^21^ as a portable, rapid, inexpensive, and highly sensitive protein binding detection platform. It introduces modular, competitive, epitope-specific binding of three proteins (ACE2, spike (or RBD), and nAb) that are of crucial importance to host cell infection, thus allowing quantification of nAb potency to block SARS-CoV-2 variants from binding to ACE2. Uniquely, NasRED utilizes ACE2 functionalized AuNPs as signal reporters, creates multivalent RBD (or full S protein) as surrogate viral antigens to modulate ACE2-AuNP clustering, accelerates the reaction by centrifugation-induced reagent localization, and eliminates the need for complex optical instruments, washing, or enzymatic reactions. With the simplified operation, digitized circuit readout, small footprint, and no need to utilize infectious virus, NasRED-nAb is an *in vitro*, in-solution, and plug-and-play assay platform suitable for semiquantitative immunity analysis with lower biosafety requirements and a POC setting. While it is not designed to replace virus-based neutralization assays (**Table S3**), it offers a rapid alternative to traditional nAb detection platforms to test target mAbs’ activity against different virus variants when the dominant virus receptor(s) is known. NasRED-nAb could distinguish the neutralizing activity level of different mAbs in PBS and serum samples of up to 4 orders of magnitude in concentration ranges (ng/mL-µg/mL), feasible for testing human serum COVID-19 IgG concentrations (e.g. 1.4 to 4200 µg/mL^39^) that may be diluted when needed, with its neutralization signals correlated with hACE2 binding competitive ELISA data. Furthermore, in longitudinal serological studies, NasRED-nAb proved effective in tracking an individual’s immunity changes in nAb levels over time. Such semiquantitative analysis could offer valuable insight into immune monitoring and therapeutic evaluation. Unlike LFA^16^, NasRED-nAb platform can be automated to streamline sample handling and data analysis, and adapted for high-throughput applications to enable broad serological testing. In addition, the assay platform could be explored in the future to analyze a broader range of biological fluids, including whole blood, to further enhance its clinical relevance.

## Materials and methods

### Sources of reagents and instrumentation

Biotinylated WT-RBD (Wuhan-Hu-1, 28.2 kDa, cat.# SPD-C82E9), WT-S1 subunit (Wuhan-Hu-1, 78.6 kDa, cat.# S1N-C82E8), Gamma-RBD (P1-20J/501Y.V3, 28.2 kDa, cat.# SPD-C82E7), Omicron-RBD (BF.7 and BA.4.6, 28.3 kDa, cat.# SPD-C82E1), ACE2 (111.7 kDa, cat.# AC2-H82F9), and neutralizing antibody AS35 (Human IgG1, cat# SAD-S35) were purchased from ACROBiosystems (see sequences in the supplementary). The WT-, Gamma-, and Omicron-RBD proteins (residues 319-537) and WT-S1 proteins (residues 16 - 685) all have a polyhistidine tag and an Avi tag (Avitag™) at the C-terminus. The ACE2 protein contains a human IgG1 Fc tag at the C-terminus, followed by an Avi tag. Biotinylation of the RBD, S1, and ACE2 proteins was performed enzymatically by the manufacturer using Avitag™ technology to tag the single lysine residue in the Avitag. The non-neutralizing antibody CR3022 (cat.# CR3022) was purchased from Absolute Antibody. The microcentrifuge (accuSpin Micro 17) and the vortexer (Analog Vortex Mixer Catalog No. 02-215-365) were both purchased from Thermo Fisher.

The murine mAbs tested in this work (SARS2-02, SARS2-03, SARS2-10, SARS2-31, SARS2-38, and SARS2-71) have been published previously^27^. Antibodies were produced in hybridoma cells and purified from supernatants by protein A affinity chromatography.

### 1×PBS dilution buffer

The 10×Phosphate-buffered saline (10×PBS, from Fisher Scientific) stock solution was combined with glycerol (from Sigma-Aldrich), BSA (from Sigma-Aldrich), and deionized water (from Fisher Scientific) to create a 1×PBS dilution buffer. This buffer had a final concentration of 1×PBS, 20% v/v glycerol, and 1% wt BSA. The resulting dilution buffer, with a pH of approximately 7.4, was utilized to prepare AuNP sensors, antibody solutions, and diluted biological media.

### Electronic readout system

An electronic readout system, referred to as PED here, similar to our previous studies^19–21^, consisted of three key components (**Fig. S1**): an LED (WP7113PGD, Kingbright), a photodiode-integrated circuit (SEN-12787, SparkFun Electronics, integrated with a digital light-sensor APDS-9960 from Broadcom), and a 3D printed (Qidi Tech X-Plus 3D Printer) black carbon fiber polycarbonate filament tube holder to fit snugly into a standard 0.5 mL Labcon microcentrifuge tube. The photodiode APDS-9960 was biased at 3.3 V and interfaced with a microcontroller (Atmega328) to convert the output into a digital signal. The electronic components were purchased from DigiKey unless specified otherwise.

### Sensing solution (RBD/S1 or ACE2-AuNP) preparation and quantification

Streptavidin-coated AuNPs (∼0.13 nM, 80 nm, optical density (OD)10) were purchased from Cytodiagnostics. Then, 50 μL of such AuNPs were mixed with an excess of biotinylated antigen (S or RBD) or ACE2 proteins (*e.g*., approximately 2 μM, 20 μL), incubated for 2 h, and diluted with 1 mL of 1×PBS dilution buffer. The mixture was purified by centrifugation at 9,600× *g* for 10 min, followed by removing 1050 μL supernatant and adding 1050 μL 1×PBS dilution buffer. This purification step was repeated two times to remove any unbound biotinylated antigen or ACE2 protein. The purified sensing solutions were measured using PED to determine the AuNP ^*OD*^*PED*, and then adjusted to the desired concentration by mixing with 1×PBS dilution buffer (e.g., ^*OD*^*PED*∼ 0.5 for NasRED-tAb, and ^*OD*^*PED* ∼ 0.8 and 0.4 for NasRED-nAb conditions 1 and 2, respectively). Before each sensing experiment, the stock sensing solutions were aliquoted into 18 μL and 12 μL in Eppendorf tubes for NasRED-tAb and NasRED-nAb, respectively.

### OD_PED_ quantification

For background normalization, the transmission signals of the PBS buffer (24 μL, in tubes) were collected on the PED device and averaged as 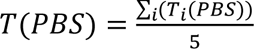 (*i*=1 to 5 for the measurements along five designated orientations). The functionalized AuNPs were again measured on the same PED device along the same five orientations and recorded as 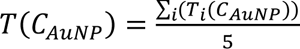 (i=1 to 5, at AuNP concentration *C*_*AuNP*_). *OD*_*PED*_ (*C*_*AuNP*_) was then calculated to correlate the AuNP concentrations with the PED measured signals following:

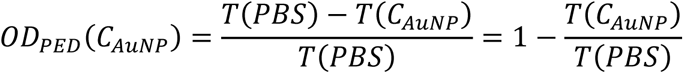

To estimate the number of AuNPs and relatively the concentration of surface proteins in the solution, we used the manufacturer’s information for OD10 AuNPs (130 pM, 670 streptavidin on the surface of the AuNPs) and followed the correlation between the manufacturer’s OD and *OD*_*PED*_ as *OD*_*PED*_ = 0.3 *OD* + 0.06 (**Fig. S7**).

### NasRED-tAb and NasRED-nAb workflow

Commercial antibodies or mAbs stock solutions (6 μM in 1×PBS) were subjected to a 10-fold serial dilution in 1×PBS dilution buffer or HPS (Innovative research) to achieve target concentrations ranging from 400 fM to 4 μM. Furthermore, the multivalent-S (or RBD) solutions were prepared by mixing streptavidin (Sigma Aldrich) and biotinylated antigens at a molar ratio of 1:3 and incubated for 2 hours. Subsequently, 6 μL of mAb in PBS or HPS solutions were mixed with 18 μL of the AuNP sensing solution for NasRED-tAb assay. Differently for the NasRED-nAb assay, the 6 μL of mAb solutions (or sera samples to be tested) were mixed with 12 μL of the AuNP sensing solution and 6 μL of the multivalent-S (or RBD) solution. This resulted in a four-fold dilution of the mAb solution or sera sample in the 24 μL mixed reaction solution, yielding final mAb concentrations from 100 fM to 1 μM or 25% diluted sera. A 24 µL negative control (NC) was also designed as a mixture of either 18 µL of functionalized AuNPs and 6 µL of tested medium (without mAbs) for NasRED-tAb, or 12 µL of such AuNPs, 6 µL of multivalent-RBD, and 6 µL of tested medium (PBS or HPS, without mAbs) for NasRED-nAb. The mixed mAb and NC reaction solutions were then vortexed at 2,050 RPM for 5 seconds for homogenized reagent distribution. For rapid detection, the assay solution was centrifuged at 3,500 RPM (1,200 g) for 5 min, incubated for 20 min, and vortexed at 2,050 RPM for 5 seconds before readout.

### NasRED data processing

The analysis of NasRED data consisted of signal calibration and normalization. The samples were measured using our PED device. Each mAb-containing sample and the NC were measured at five designed orientations to account for variations in tube alignment. The digital photodiode sensor readings from each sample were averaged and stored in datasheets (.csv), with all data collection and storage automated via Python scripts. The averaged signal 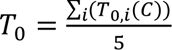 where i = 1 to 5 for the five measurements at antibody concentration C, represented the overall optical transmission through the sample and the microcentrifuge tube measured immediately after mixing. Following standard procedures, including centrifugation, incubation, and vortex agitation, another set of five measurements was taken for each sample along the same orientations, and the signals were averaged again to give 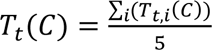 (*i*=1 to 5). The calibrated signals, reflecting the change in optical transmission due to AuNP precipitation, were obtained as *T_D,i_*(*C*) = *T_t,i_* (*C*) – *T*_0,*i*_ (*C*) and averaged as 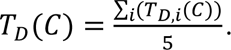 The NC signal difference *T_D_* (*NC*) = *T*_*t*_(*NC*) − *T*_0_(*NC*) was also computed, and *T*_*D*_(*NC*) <5% *T*_0_(*NC*) was used as a criterion to confirm that most AuNPs returned to their original state (redispersion in the case of NasRED-tAb) after mixing. For Nasred-nAb, *T*_*t*_(*NC*) ∼ *T*_*t*_(*PBS or Sera*) was used as a measure of confirmation that almost all AuNPs precipitate in the absence of nAbs.

A positive control (PC) was introduced for normalization, representing complete AuNP precipitation and maximum optical transmission in NasRED-tAb and complete AuNP redispersion for NasRED-nAb. For NasREd-tAb, 24 μL 1×PBS dilution buffer was used, and its transmission was recorded as the PC signal *T*(*PC*) = *T*_*t*_(*PBS*). For NasRED-nAb, the PC reference was 12 μL of ACE2-AuNPs mixed with 6 μL of multivalent-RBD (or S1) in the designated conditions (2 nM and 1 nM for conditions 1 and 2, respectively) and 6 μL SARS2-38 spiked in either PBS or HPS at a high concentration (*C*_*H*_=10 μM). Here, the PC in NasRED-nAb acted as a concentration reference for the unknown sera samples as well and was defined as *T*(*PC*) = *T*_*t*_(*C*_*H*_) − *T*_0_(*C*_*H*_) = *T*_*D*_(*C*_*H*_).

To account for small differences in sensor concentration between batches and the significant variations in biological media, the NasRED signals were subsequently normalized on a scale from 0 to 1, calculated as transmission signal 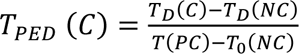 for NasRED-tAb or an extinction signal 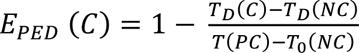 for NasREd-nAb. Additionally, the measurement errors were calculated as the standard deviation (SD) of the five normalized signals, 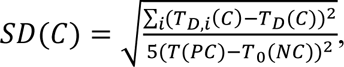 with *i* representing the five orientations. The normalized sensing signals *T*_*PED*_(*C*) ± *SD*(*C*) *or E*_*PED*_(*C*) ± *SD*(*C*) were plotted against the mAb concentrations.

### Data Fitting and LoD Calculations

The normalized mAb signals *S*_*PED*_(*C*) ( *T*_*PED*_(*C*) for NasRED-tAb and *E*_*PED*_(*C*) for NasRED-nAb) were fitted using the orthogonal distance regression algorithm in Origin 2024b software (OriginLab, USA), which considers both the data and error weight in the fitting calculations. A dose-response model (sigmoidal function) was used for fitting:

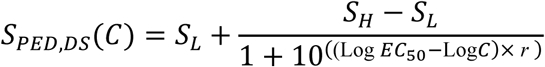

Where *S*_*H*_and *S*_*L*_are the highest and lowest sensing signals observed, EC50 is the half-maximal concentration, and *r* determines the slope of the curve. A large signal contrast *ΔS*_*PED*_ = *S*_*H*_ − *S*_*L*_, in combination with a shallow slope *r*, is desirable to achieve a wide dynamic range. The fitting parameters were adjusted to optimize goodness-of-fit metrics such as reduced Chi-square, R-squared, and reduced sum of squares (RSS), aiming for values near 1, 1, and 0, respectively. The limit of detection (LoD) for NasRED-tAb was determined by finding the lowest concentration where the PED signal significantly differs from the background, following *S*_*PED*,*DS*_(*LoD*) = *S*_*PED*_(*NC*) + 1.645 × (*SD*(*NC*) + *SD*(*C*_*Min*_)) where *C*_*Min*_is the lowest protein concentration tested^40,41^.

### Human Sera Samples and ethical approval

Human serum samples used in this study were collected from two serosurvey studies conducted in September 2021 and October 2022. Both studies involving human participants were approved by the Arizona State University IRB (ID: STUDY00014505 and STUDY00015522). Participants provided informed consent prior to participating in these studies.

## Acknowledgements

This project was partly supported by the National Institutes of Health under grant no. R21AI169098, R01 AI157155, and DP2GM149552, by the U.S. Department of Agriculture under grant no. AFRI 2022-67021-37013, and by the National Science Foundation under grant no. 1847324. C. Wang acknowledges Dr. Qin Xu at NIH for helpful suggestions of protein products and Dr. Neal Woodbury for critical reading of the draft.

## Supplementary

### 1. NasRED readout

**Figure S1.**
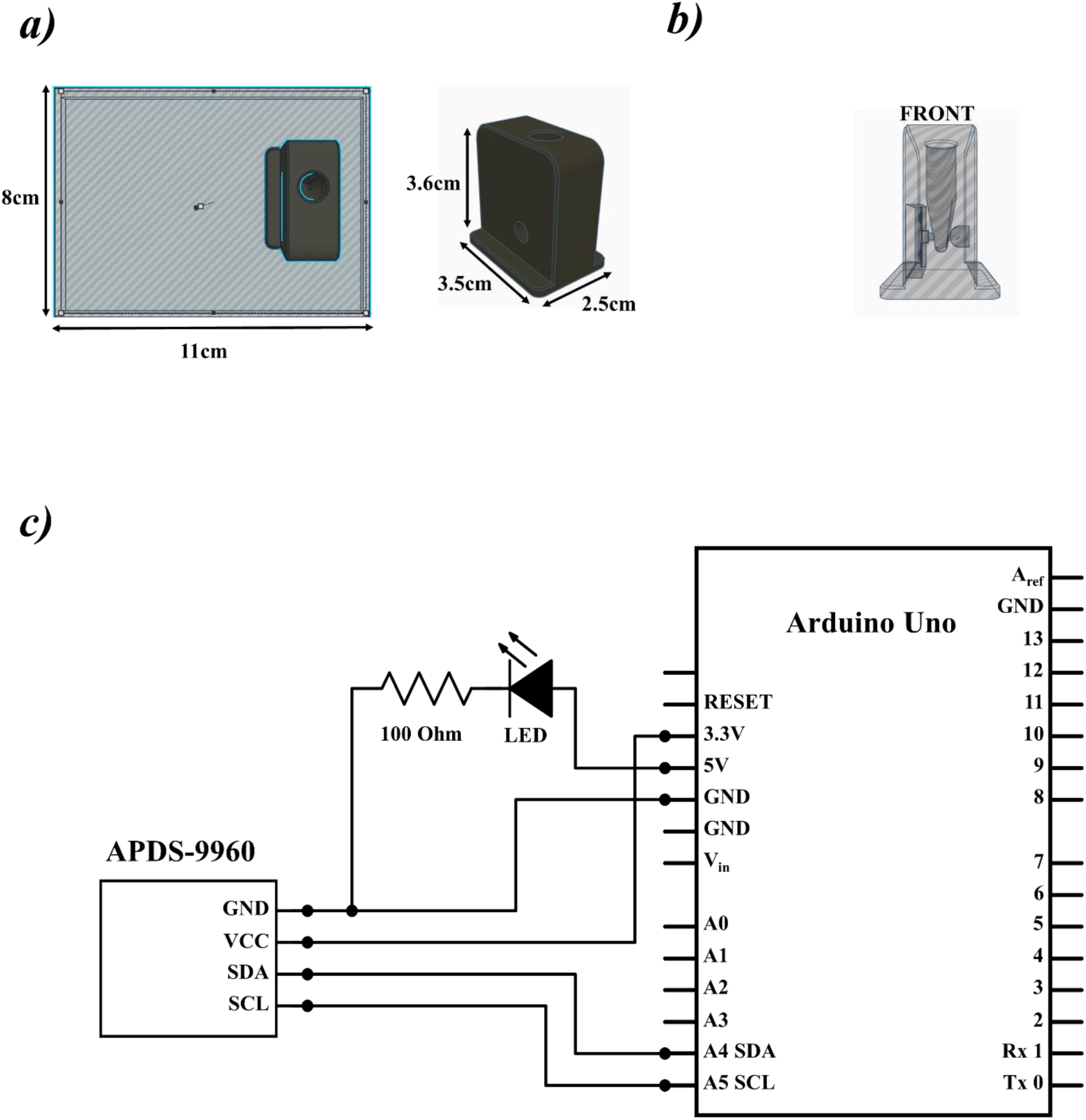
Portable Electronic Readout (PED) schematics. (a) CAD design of the microcentrifuge 3D printed device box tube holder and (b) the inner perspective of the light path, sensor, and LED locations placed at the upper solution level to avoid aggregate light interactions. (c) A custom circuit design based on the commercial Arduino Uno, comprising a voltage regulator, is used to power the LED and input and output ports for signal digitalization using a computer.

### 2. NasRED-tAb protocol and extended data

**Figure S2.**
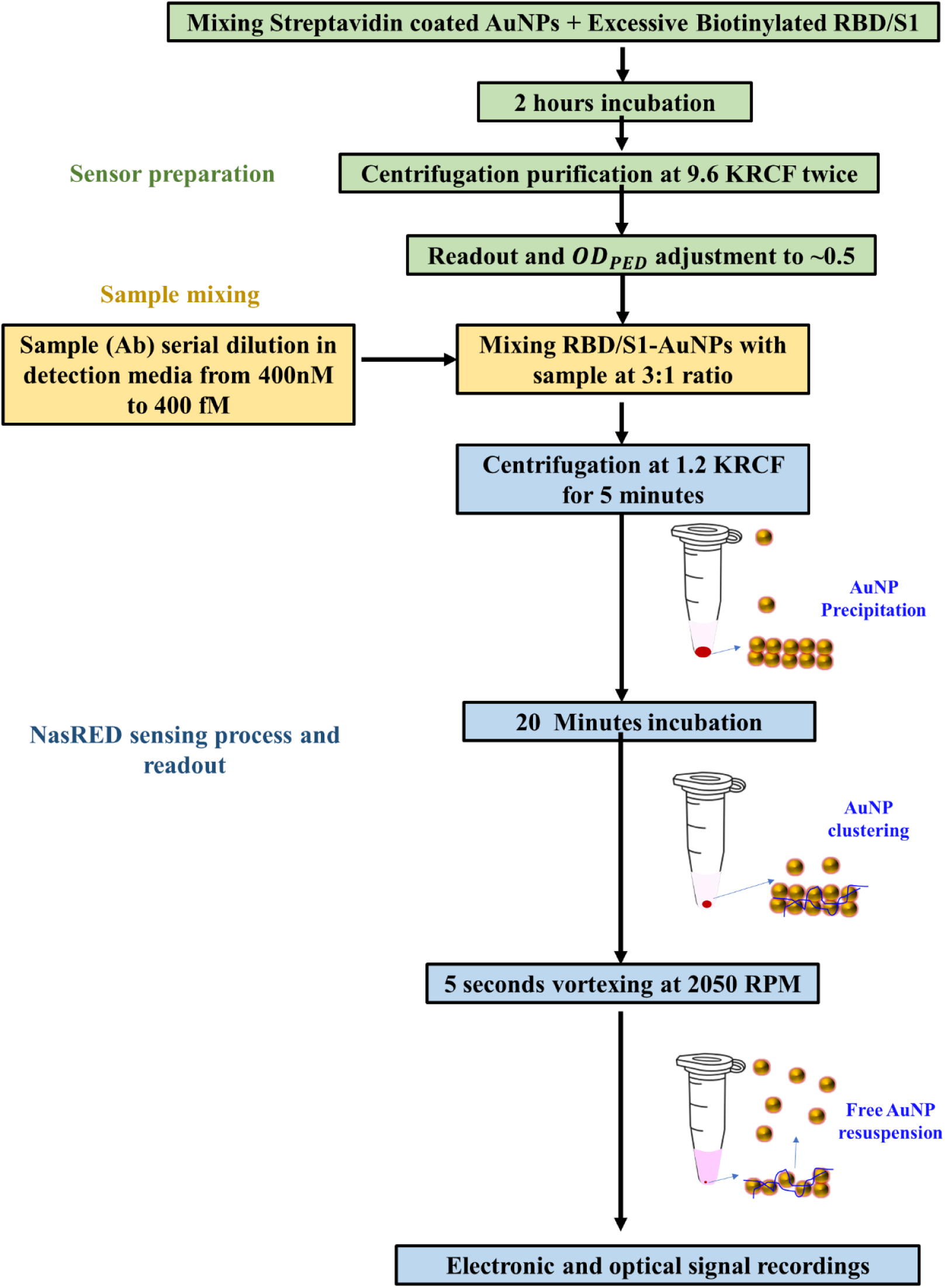
NasRED-tAb Workflow.

**Figure S3.**
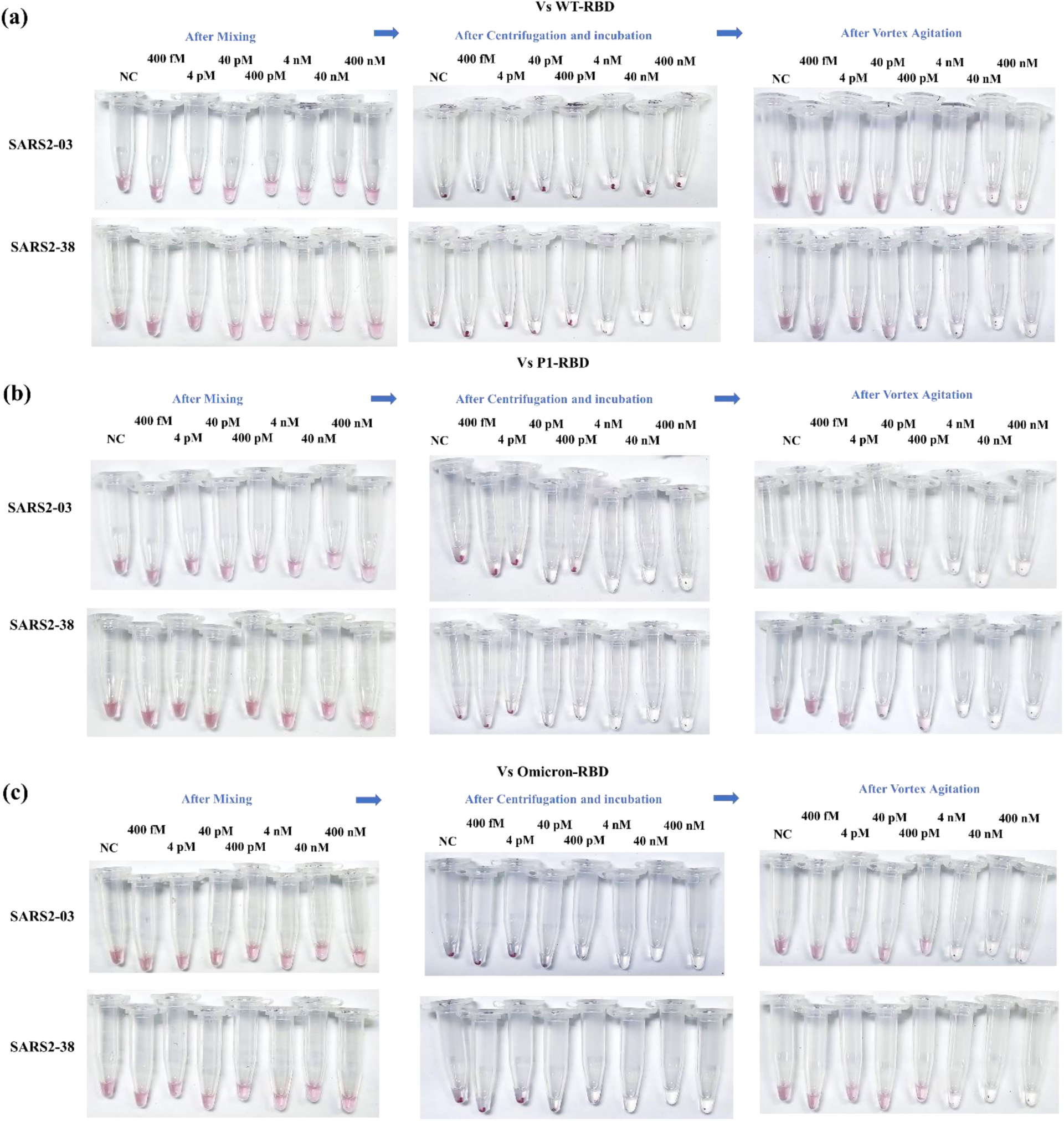
Optical images of NasRED-tab test tubes at different stages during the sensing procedure. (a-c) Test tubes for mAbs (SARS2-03 and SARS2-38) against: (a) WT-RBD, (b) P1-RBD, and (c) Omicron-RBD. Left column: after mixing; middle column: after centrifugation and incubation; and right column: after vortex agitation.

### 3. NasRED-nAb protocol and extended data

**Figure S4.**
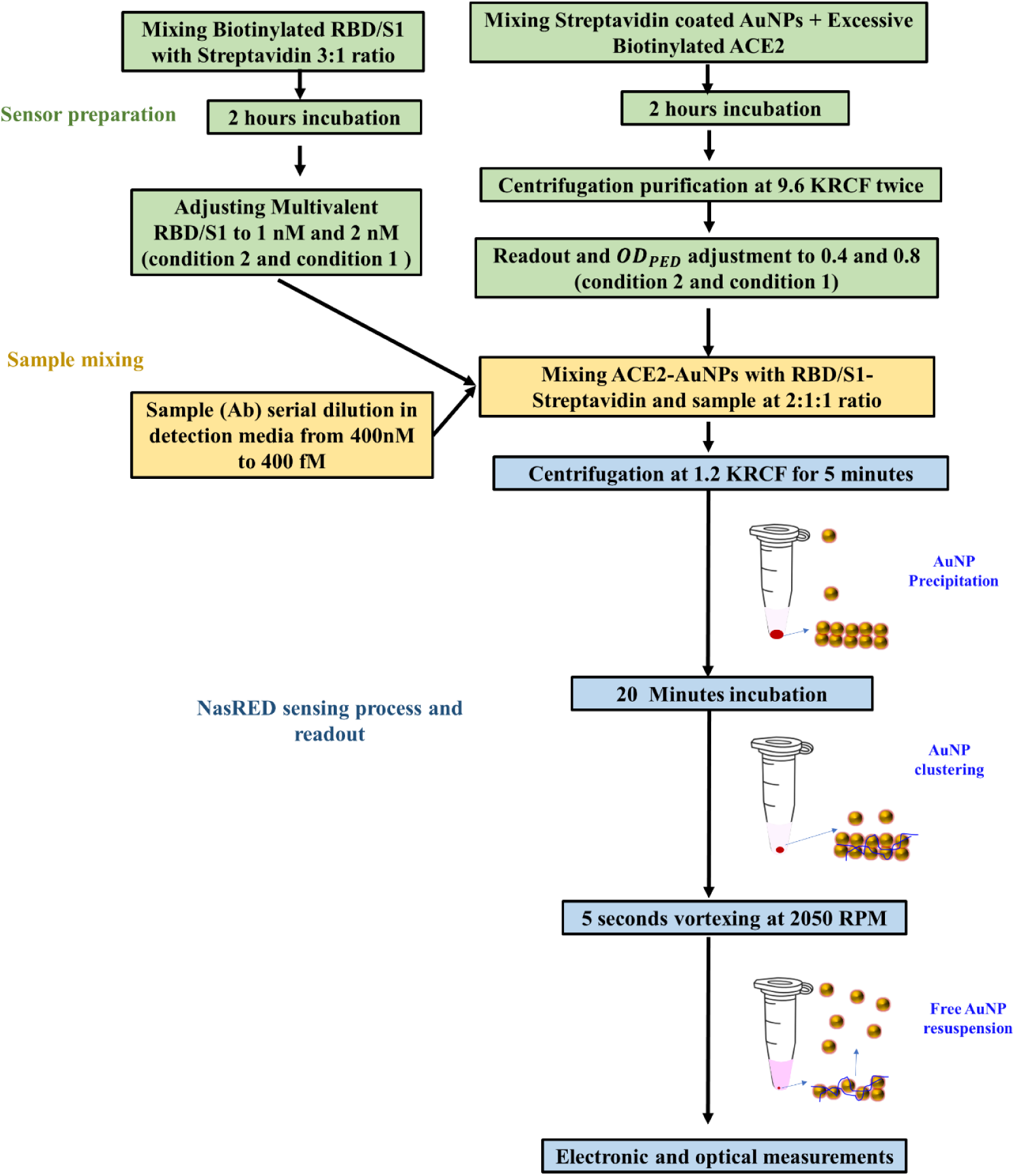
NasRED-nAb Workflow.

**Figure S5.**
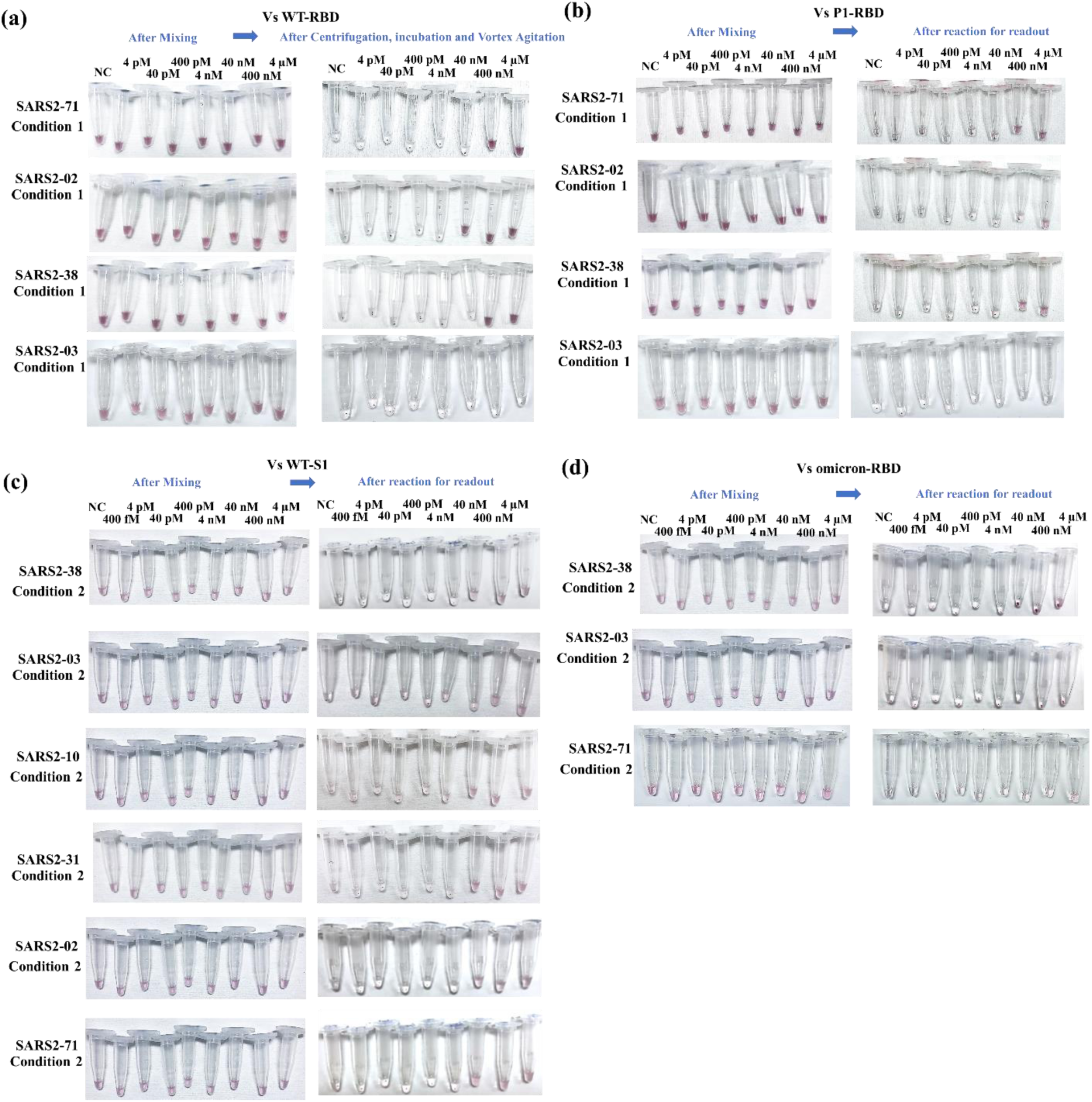
Optical images of NasRED-nab test tubes during the sensing procedure. (a) Test tubes of six mAbs (SARS2-02, -03, -10, -31, -38, and -71) against: (a) WT-RBD, (b) P1-RBD, (c) WT-S1, and (d) Omicron-RBD antigens. Left column: after mixing; right column: after reaction for readout.

**Figure S6.**
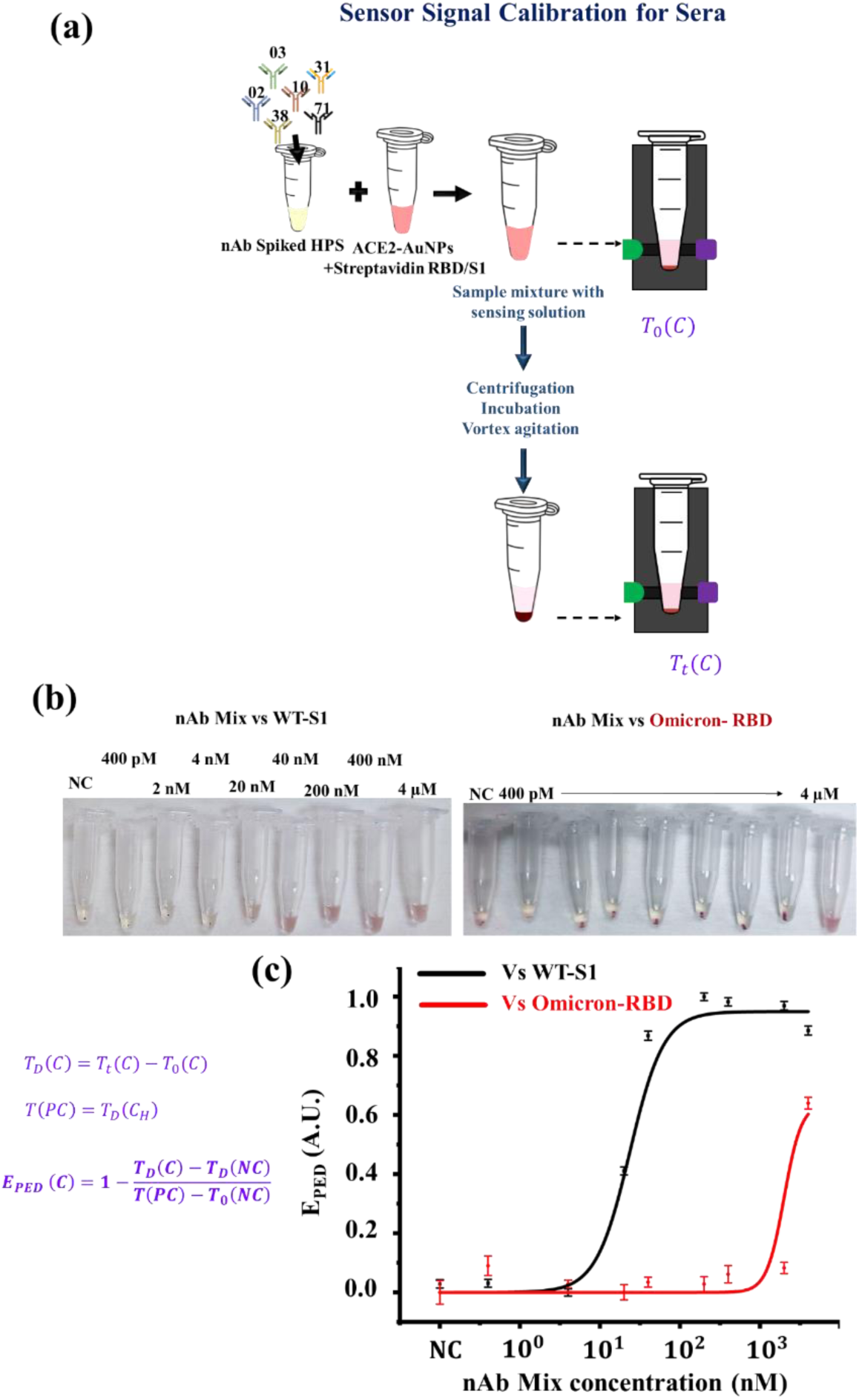
NasRED-nAb sensor calibration utilizing mixed mAbs spiked in Human pooled serum (HPS). (a) schematics of the analysis and calibration steps for sera samples. (b) Optical images of mAb mixture of SARS2-02, SARS2-03, SARS2-10, SARS2-31, SARS2-38, and SARS2-71 (equal ratio) spiked in human pooled serum at different concentrations against diluted multivalent-WTS1 (left) and Omicron-RBD (right). The yellowish serum color is visible in the negative control (NC) sample (no mAb). (c) Normalized optical extinction signals *E*_*PED*_ plotted against the mAb concentrations versus Omicron-RBD (red) and WT-S1 (black). Here, the centrifugation for sera samples was 1,600 RCF, and the vortexing speed was 2,250 RPM. The background light absorbance of the sera samples (yellowish color) was removed during calibration, where the transmission signal of the tubes after the reaction *T*_*t*_(C) was subtracted from the transmission of the same tubes after mixing reagents *T*_0_(C). The resultant data *T*_*D*_(C) represented the changes in the optical transmission only caused by the AuNPs.

### 4. *OD_PED_* correlation with manufacturer-provided OD data

**Figure S7.**
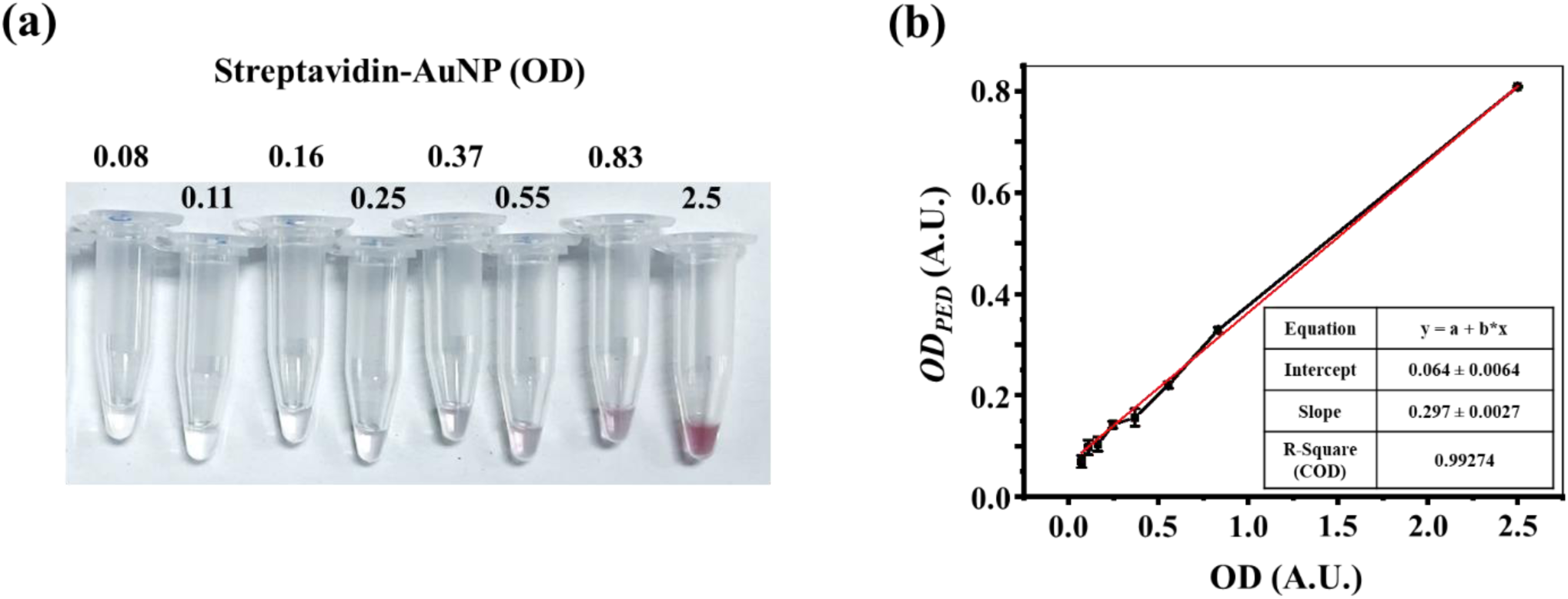
Correlation of *OD_PED_* with the manufacturer’s OD. (a) Images of tubes displaying color changes in AuNP solutions of different concentrations corresponding to the optical density (OD) values provided by the manufacturer (Cytodiagnostics). The AuNPs obtained from Cytodiagnostics were reported to have an initial optical density OD of 10. In this context, OD is the base-10 logarithm of the ratio between incident and transmitted optical power. A higher OD indicates greater extinction and a higher concentration of AuNPs. (b) The electronic readout signal *OD*_*PED*_ plotted as a function of the manufacturer-provided OD values, with a linear fit described by the equation *OD*_*PED*_ = 0.3 × *OD* + 0.06.

### 5. Sera sample information

**Table S1.**
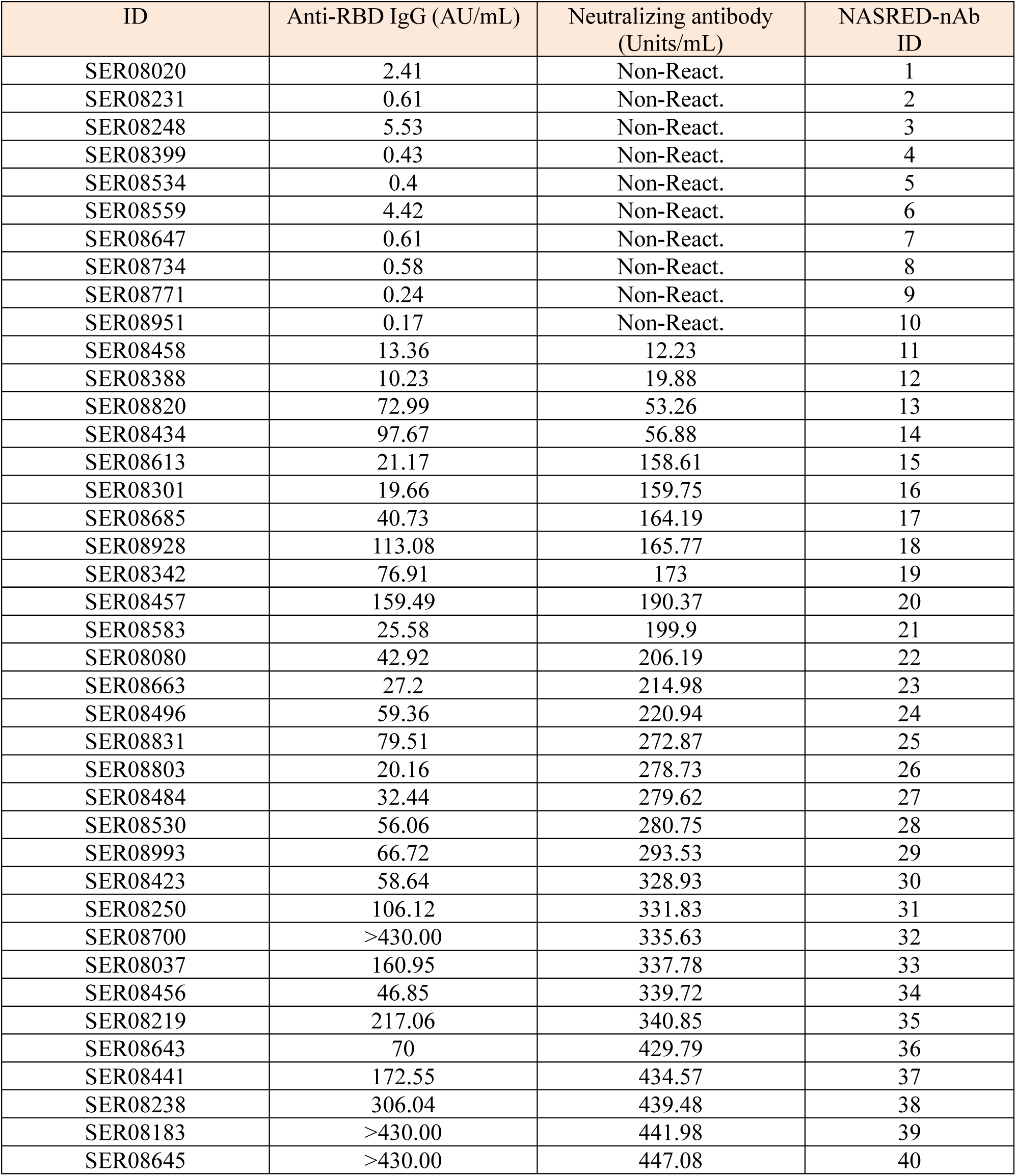
Randomly selected sera samples from Serosurvey 1 (September 2021) and corresponding Total antibody and hACE2 binding competitive ELISA assay results.

| ID | Anti-RBD IgG (AU/mL) | Neutralizing antibody (Units/mL) | NASRED-nAb ID |
| --- | --- | --- | --- |
| SER08020 | 2.41 | Non-React. | 1 |
| SER08231 | 0.61 | Non-React. | 2 |
| SER08248 | 5.53 | Non-React. | 3 |
| SER08399 | 0.43 | Non-React. | 4 |
| SER08534 | 0.4 | Non-React. | 5 |
| SER08559 | 4.42 | Non-React. | 6 |
| SER08647 | 0.61 | Non-React. | 7 |
| SER08734 | 0.58 | Non-React. | 8 |
| SER08771 | 0.24 | Non-React. | 9 |
| SER08951 | 0.17 | Non-React. | 10 |
| SER08458 | 13.36 | 12.23 | 11 |
| SER08388 | 10.23 | 19.88 | 12 |
| SER08820 | 72.99 | 53.26 | 13 |
| SER08434 | 97.67 | 56.88 | 14 |
| SER08613 | 21.17 | 158.61 | 15 |
| SER08301 | 19.66 | 159.75 | 16 |
| SER08685 | 40.73 | 164.19 | 17 |
| SER08928 | 113.08 | 165.77 | 18 |
| SER08342 | 76.91 | 173 | 19 |
| SER08457 | 159.49 | 190.37 | 20 |
| SER08583 | 25.58 | 199.9 | 21 |
| SER08080 | 42.92 | 206.19 | 22 |
| SER08663 | 27.2 | 214.98 | 23 |
| SER08496 | 59.36 | 220.94 | 24 |
| SER08831 | 79.51 | 272.87 | 25 |
| SER08803 | 20.16 | 278.73 | 26 |
| SER08484 | 32.44 | 279.62 | 27 |
| SER08530 | 56.06 | 280.75 | 28 |
| SER08993 | 66.72 | 293.53 | 29 |
| SER08423 | 58.64 | 328.93 | 30 |
| SER08250 | 106.12 | 331.83 | 31 |
| SER08700 | >430.00 | 335.63 | 32 |
| SER08037 | 160.95 | 337.78 | 33 |
| SER08456 | 46.85 | 339.72 | 34 |
| SER08219 | 217.06 | 340.85 | 35 |
| SER08643 | 70 | 429.79 | 36 |
| SER08441 | 172.55 | 434.57 | 37 |
| SER08238 | 306.04 | 439.48 | 38 |
| SER08183 | >430.00 | 441.98 | 39 |
| SER08645 | >430.00 | 447.08 | 40 |

**Table S2.** Sera samples information for the longitudinal study. Samples were collected from the same individuals with identical conditions in September 2021 in Serosurvey 1 when all subjects had been vaccinated against the wildtype variant (Wuhan-Hu-1) and in October 2022 in Serosurvey 2 after the same individuals’ infection by the Omicron Variant (BF.7 & and BA.4.6). Pseudo ID has been assigned to individuals and Serum aliquot number for collected samples.

| Collection Time | 09/2021 | 10/2022 |
| --- | --- | --- |
| Subject Condition | Post Vaccination Vs WT | Post Infection by Omicron |
| Survey Type | Sero1 | Sero2 |
| Pseudo ID | Serum Aliquot # | Serum Aliquot # |
| 1 | SER08810 | SST100513 |
| 23 | SER08161 | SST100351 |
| 30 | SER08949 | SST100873 |
| 42 | SER08681 | SST100619 |
| 53 | SER08568 | SST101381 |
| 67 | SER08717 | SST100497 |
| 71 | SER08164 | SST100976 |
| 100 | SER08213 | SST100167 |
| 111 | SER08396 | SST100752 |
| 133 | SER08569 | SST101303 |

### 6. Technology Comparison

**Table S3.** Summary of neutralization test technologies.

| Assay Format | Epitope Specificity | Instrument/ environment | Antibody quantification | Assay time | Reference |
| --- | --- | --- | --- | --- | --- |
| Plaque/Foci Reduction Neutralization Assay | Very high (live virus) | Stringent (BSL3) | Quantitative (LoD ~ nM) | 2-7days | <sup>1,2</sup> |
| Micro Neutralization Assay | Very high (live virus) | Stringent (BSL3) | Quantitative (LoD ~ nM) | 2 days | <sup>1,3,4</sup> |
| Pseudo virus Neutralization Assay | High (pseudo virus) | Accessible (BSL2) | Quantitative (LoD ~ nM) | 2.5 days | <sup>1,5</sup> |
| Surrogate Neutralizing Assay | High (viral proteins) | Accessible (BSL1 or2) | Quantitative (LoD ~ nM) | 1-2 hours | <sup>6,7</sup> |
| hACE2 binding competitive ELISA Assay | High (viral proteins) | Accessible (BSL2) | Quantitative (LoD ~ nM) | 1-3 hours | <sup>8</sup> |
| AVNIR™ Technology | High (viral proteins) | Accessible (BSL2) | Quantitative (LoD ~ nM) | 1-2 days | <sup>9</sup> |
| Microfluidic Antibody profiling | High (viral proteins) | Accessible (BSL2) | Quantitative (LoD ~ nM) | 1-2 hours | <sup>10</sup> |
| Competitive Chemiluminescence Immunoassay | High (viral proteins) | Accessible (BSL1 or2) | Semiquantitative | 30-40 minutes | <sup>11</sup> |
| Bead-based ACE2-RBD inhibition assay | High (viral proteins) | Accessible (BSL1 or2) | Quantitative (LoD ~ nM) | 4 hours | <sup>12</sup> |
| Lab-on-a-chip Concurrent Electrochemical detection | High (viral proteins) | Accessible (BSL1 or2) | Quantitative (LoD ~ nM) | 1-2 hours | <sup>13</sup> |
| Double-antigen Sandwich Lateral Flow Immunoassay | High (viral proteins) | Accessible (BSL1 or2) | Quantitative (LoD ~ nM) | 20-30 minutes | <sup>14-17</sup> |
| Semiquantitative Paper-Based Microfluidic | High (viral proteins) | Accessible (BSL1 or2) | Semiquantitative | 15 minutes | <sup>18</sup> |
| Upconversion nanoparticle-based neutralizing immunoassay (UNIK) | High (viral proteins) | Accessible (BSL1 or2) | Quantitative (LoD ~ nM) | 1 hour | <sup>19</sup> |
| Blockade of ACE-2 binding (BoAb) assay | High (viral proteins) | Accessible (BSL 2) | Quantitative (LoD ~ nM) | 4 hours | <sup>20</sup> |
| Polymer Brush Surface Microarrays | High (viral proteins) | Accessible (BSL1 or2) | Quantitative (LoD ~ nM) | 30 minutes | <sup>21</sup> |
| Quantitative microfluidic assay | High (viral proteins) | Accessible (BSL1 or2) | Quantitative (LoD ~ nM) | 5-10 minutes | <sup>22</sup> |
| MetaSPR VNT biosensor | High (viral proteins) | Accessible (BSL1 or2) | Quantitative (LoD ~ fM) | 20 minutes | <sup>23</sup> |
| Flow cytometry-based neutralization assay | High (viral proteins) | Accessible (BSL1 or2) | Quantitative (LoD ~ nM) | 60-90 minutes | <sup>24</sup> |
| Electrochemical Immunosensor | High (viral proteins) | Accessible (BSL1 or2) | Quantitative (LoD ~ fM) | 15-25 minutes | <sup>25,26</sup> |
| NasRED-tAb and NasRED-nAb | High (viral proteins) | Accessible (BSL1 or2) | Quantitative (LoD ~ fM) | 20-30 minutes | This study |

### 7. Protein sequences

#### WT-RBD

rvqptesivrfpnitnlcpfgevfnatrfasvyawnrkrisncvadysvlynsasfstfkcygvsptklndlcftnvyadsfvirgdevrqiapgqtgkiadynyklpddftgcviawnsnnldskvggnynylyrlfrksnlkpferdisteiyqagstpcngvegfncyfplqsygfqptngvgyqpyrvvvlsfellhapatvcgpkkstnlvknk—Linker(gggsgggs)—10*His(hhhhhhhhhh)—Avi(15AA)

#### WT-S1

vnlttrtqlppaytnsftrgvyypdkvfrssvlhstqdlflpffsnvtwfhaihvsgtngtkrfdnpvlpfndgvyfasteksniirgwifgttldsktqsllivnnatnvvikvcefqfcndpflgvyyhknnkswmesefrvyssannctfeyvsqpflmdlegkqgnfknlrefvfknidgyfkiyskhtpinlvrdlpqgfsaleplvdlpiginitrfqtllalhrsyltpgdsssgwtagaaayyvgylqprtfllkynengtitdavdcaldplsetkctlksftvekgiyqtsnfrvqptesivrfpnitnlcpfgevfnatrfasvyawnrkrisncvadysvlynsasfstfkcygvsptklndlcftnvyadsfvirgdevrqiapgqtgkiadynyklpddftgcviawnsnnldskvggnynylyrlfrksnlkpferdisteiyqagstpcngvegfncyfplqsygfqptngvgyqpyrvvvlsfellhapatvcgpkkstnlvknkcvnfnfngltgtgvltesnkkflpfqqfgrdiadttdavrdpqtleilditpcsfggvsvitpgtntsnqvavlyqdvnctevpvaihadqltptwrvystgsnvfqtragcligaehvnnsyecdipigagicasyqtqtnsprrar—Linker(gggsgggs)—10*His(hhhhhhhhhh)—Avi(15AA)

#### Gamma(P1)-RBD

rvqptesivrfpnitnlcpfgevfnatrfasvyawnrkrisncvadysvlynsasfstfkcygvsptklndlcftnvyadsfvirgdevrqiapgqtgniadynyklpddftgcviawnsnnldskvggnynylyrlfrksnlkpferdisteiyqagstpcngvkgfncyfplqsygfqptygvgyqpyrvvvlsfellhapatvcgpkkstnlvknk—Linker(gggsgggs)—10*His(hhhhhhhhhh)—Avi(15AA)

#### Omicron-RBD

rvqptesivrfpnitnlcpfdevfnattfasvyawnrkrisncvadysvlynfapffafkcygvsptklndlcftnvyadsfvirgnevsqiapgqtgniadynyklpddftgcviawnsnkldskvggnynyryrlfrksnlkpferdisteiyqagnkpcngvagvncyfplqsygfrptygvghqpyrvvvlsfellhapatvcgpkkstnlvknk—Linker(gggsgggs)—10*His(hhhhhhhhhh)—Avi(15AA)

#### ACE-2

qstieeqaktfldkfnheaedlfyqsslaswnyntniteenvqnmnnagdkwsaflkeqstlaqmyplqeiqnltvklqlqalqqngssvlsedkskrlntilntmstiystgkvcnpdnpqeclllepglneimansldynerlwaweswrsevgkqlrplyeeyvvlknemaranhyedygdywrgdyevngvdgydysrgqliedvehtfeeikplyehlhayvraklmnaypsyispigclpahllgdmwgrfwtnlysltvpfgqkpnidvtdamvdqawdaqrifkeaekffvsvglpnmtqgfwensmltdpgnvqkavchptawdlgkgdfrilmctkvtmddfltahhemghiqydmayaaqpfllrnganegfheavgeimslsaatpkhlksigllspdfqedneteinfllkqaltivgtlpftymlekwrwmvfkgeipkdqwmkkwwemkreivgvvepvphdetycdpaslfhvsndysfiryytrtlyqfqfqealcqaakhegplhkcdisnsteagqklfnmlrlgksepwtlalenvvgaknmnvrpllnyfeplftwlkdqnknsfvgwstdwspyadqsikvrislksalgdkayewndnemylfrssvayamrqyflkvknqmilfgeedvrvanlkprisfnffvtapknvsdiiprtevekairmsrsrindafrlndnsleflgiqptlgppnqppvs——Linker(gggsgggs)——Fc(pkssdkthtcppcpapellggpsvflfppkpkdtlmisrtpevtcvvvdvshedpevkfnwyvdgvevhnaktkpreeqynstyrvvsvltvlhqdwlngkeykckvsnkalpapiektiskakgqprepqvytlppsrdeltknqvsltclvkgfypsdiavewesngqpennykttppvldsdgsfflyskltvdksrwqqgnvfscsvmhealhnhytqkslslspgk)——Avi(15AA)

## References

1. COVID-19 CORONAVIRUS PANDEMIC. https://www.worldometers.info/coronavirus/.

2. Wang, Y., Wang, L., Cao, H. & Liu, C. SARS-CoV-2 S1 is superior to the RBD as a COVID-19 subunit vaccine antigen. J Med Virol 93, 892–898 (2021).

3. Schmitz, A. J. et al. A vaccine-induced public antibody protects against SARS-CoV-2 and emerging variants. Immunity 54, 2159–2166.e6 (2021).

4. Tortorici, M. A. et al. Ultrapotent human antibodies protect against SARS-CoV-2 challenge via multiple mechanisms. Science (1979) 370, 950–957 (2020).

5. Chonira, V. et al. A potent and broad neutralization of SARS-CoV-2 variants of concern by DARPins. Nat Chem Biol 19, 284–291 (2023).

6. Peng, L. et al. Monospecific and bispecific monoclonal SARS-CoV-2 neutralizing antibodies that maintain potency against B.1.617. Nat Commun 13, 1638 (2022).

7. Cameroni, E. et al. Broadly neutralizing antibodies overcome SARS-CoV-2 Omicron antigenic shift. Nature 602, 664–670 (2022).

8. Zhang, Y., et al. An updated review of SARS-CoV-2 detection methods in the context of a novel coronavirus pandemic. Bioeng Transl Med 8, (2023).

9. Addetia, A. et al. Neutralization, effector function and immune imprinting of Omicron variants. Nature 621, 592–601 (2023).

10. VanBlargan, L. A. et al. An infectious SARS-CoV-2 B.1.1.529 Omicron virus escapes neutralization by therapeutic monoclonal antibodies. Nat Med 28, 490–495 (2022).

11. Diamond, M. S. & Kanneganti, T.-D. Innate immunity: the first line of defense against SARS-CoV-2. Nat Immunol 23, 165–176 (2022).

12. Münsterkötter, L. et al. Comparison of the Anti-SARS-CoV-2 Surrogate Neutralization Assays by TECOmedical and DiaPROPH-Med with Samples from Vaccinated and Infected Individuals. Viruses 14, 315 (2022).

13. Bewley, K. R. et al. Quantification of SARS-CoV-2 neutralizing antibody by wildtype plaque reduction neutralization, microneutralization and pseudotyped virus neutralization assays. Nat Protoc 16, 3114–3140 (2021).

14. Winichakoon, P. et al. Diagnostic performance between in-house and commercial SARS-CoV-2 serological immunoassays including binding-specific antibody and surrogate virus neutralization test (sVNT). Sci Rep 13, (2023).

15. Klüpfel, J. et al. Fully Automated Chemiluminescence Microarray Analysis Platform for Rapid and Multiplexed SARS-CoV-2 Serodiagnostics. Anal Chem 94, 2855–2864 (2022).

16. Zhang, Y. et al. Development of receptor binding domain-based double-antigen sandwich lateral flow immunoassay for the detection and evaluation of SARS-CoV-2 neutralizing antibody in clinical sera samples compared with the conventional virus neutralization test. Talanta 255, 124200 (2023).

17. Najjar, D. et al. A lab-on-a-chip for the concurrent electrochemical detection of SARS-CoV-2 RNA and anti-SARS-CoV-2 antibodies in saliva and plasma. Nat Biomed Eng 6, 968–978 (2022).

18. Rahmati, Z., Roushani, M., Hosseini, H. & Choobin, H. An electrochemical immunosensor using SARS-CoV-2 spike protein-nickel hydroxide nanoparticles bio-conjugate modified SPCE for ultrasensitive detection of SARS-CoV-2 antibodies. Microchemical Journal 170, 106718 (2021).

19. Ikbal, M. D. A., Kang, S., Chen, X., Gu, L. & Wang, C. Picomolar-Level Sensing of Cannabidiol by Metal Nanoparticles Functionalized with Chemically Induced Dimerization Binders. ACS Sens 8, 4696–4706 (2023).

20. Chen, X. et al. Synthetic nanobody-functionalized nanoparticles for accelerated development of rapid, accessible detection of viral antigens. Biosens Bioelectron 202, 113971 (2022).

21. Choi, Y. et al. Nanoparticle-Supported, Rapid, and Electronic Detection of SARS-CoV-2 Antibodies and Antigens at Attomolar Level. bioRxiv 2024.09.04.611305 (2024) preprint at doi:10.1101/2024.09.04.611305.

22. Mirjalili, S., Tohidi Moghadam, T. & H. Sajedi, R. Facile and Rapid Detection of Microalbuminuria by Antibody-Functionalized Gold Nanorods. Plasmonics 17, 1269–1277 (2022).

23. Yi, C. et al. Comprehensive mapping of binding hot spots of SARS-CoV-2 RBD-specific neutralizing antibodies for tracking immune escape variants. Genome Med 13, 164 (2021).

24. Wu, N. C. et al. A natural mutation between SARS-CoV-2 and SARS-CoV determines neutralization by a cross-reactive antibody. PLoS Pathog 16, e1009089 (2020).

25. Zahmatkesh, S., Sillanpaa, M., Rezakhani, Y. & Wang, C. Review of concerned SARS-CoV-2 variants like Alpha (B.1.1.7), Beta (B.1.351), Gamma (P.1), Delta (B.1.617.2), and Omicron (B.1.1.529), as well as novel methods for reducing and inactivating SARS-CoV-2 mutants in wastewater treatment facilities. Journal of Hazardous Materials Advances 7, 100140 (2022).

26. Qu, P. et al. Enhanced neutralization resistance of SARS-CoV-2 Omicron subvariants BQ.1, BQ.1.1, BA.4.6, BF.7, and BA.2.75.2. Cell Host Microbe 31, 9-17.e3 (2023).

27. VanBlargan, L. A. et al. A potently neutralizing SARS-CoV-2 antibody inhibits variants of concern by utilizing unique binding residues in a highly conserved epitope. Immunity 54, 2399–2416.e6 (2021).

28. Bradley, Z. & Bhalla, N. Combating Prozone Effects and Predicting the Dynamic Range of Naked-Eye Nanoplasmonic Biosensors through Capture Bioentity Optimization. ACS Measurement Science Au (2024).

29. Wang, C, & Ikbal. M. D. A. Compositions and methods for rapid covid-19 detection. U.S. patent application 63116953, 17532969, (2021).

30. Yuan, M. et al. A highly conserved cryptic epitope in the receptor binding domains of SARS-CoV-2 and SARS-CoV. Science (1979) 368, 630–633 (2020).

31. Hornick, C. L. & Karush, F. Antibody affinity—III the role of multivalence. Immunochemistry 9, 325–340 (1972).

32. Zhang, J. et al. Structural and functional characteristics of the SARS-CoV-2 Omicron subvariant BA.2 spike protein. Nat Struct Mol Biol 30, 980–990 (2023).

33. Tian, X. et al. Potent binding of 2019 novel coronavirus spike protein by a SARS coronavirus-specific human monoclonal antibody. Emerg Microbes Infect 9, 382–385 (2020).

34. Fiedler, S. et al. Antibody Affinity Governs the Inhibition of SARS-CoV-2 Spike/ACE2 Binding in Patient Serum. ACS Infect Dis 7, 2362–2369 (2021).

35. Yamamoto, S. et al. Omicron BA.1 neutralizing antibody response following Delta breakthrough infection compared with booster vaccination of BNT162b2. BMC Infect Dis 23, 282 (2023).

36. Quandt, J., et al. Omicron BA.1 breakthrough infection drives cross-variant neutralization and memory B cell formation against conserved epitopes. Sci Immunol 7, (2022).

37. MacMullan, M. A. et al. ELISA detection of SARS-CoV-2 antibodies in saliva. Sci Rep 10, 20818 (2020).

38. Greenspan, NeilS. & Cooper, Laurence J. N. Cooperative binding by mouse IgG3 antibodies: implications for functional affinity, effector function, and isotype restriction. Springer Semin Immunopathol 15, (1993).

39. Song, D. et al. Rapid and quantitative detection of SARS-CoV-2 IgG antibody in serum using optofluidic point-of-care testing fluorescence biosensor. Talanta 235, 122800 (2021).

40. Tholen, D. W. et al. Protocols for determination of limits of detection and limits of quantitation; approved guideline. CLSI EP17-A 24, 34 (2004).

41. Armbruster, D. A. & Pry, T. Limit of blank, limit of detection and limit of quantitation. Clin Biochem Rev 29 Suppl 1, S49–52 (2008).

## Supplementary References

1. Bewley, K. R. et al. Quantification of SARS-CoV-2 neutralizing antibody by wildtype plaque reduction neutralization, microneutralization and pseudotyped virus neutralization assays. Nat Protoc 16, 3114–3140 (2021).

2. Horbach, I. S. et al. Plaque Reduction Neutralization Test (PRNT) Accuracy in Evaluating Humoral Immune Response to SARS-CoV-2. Diseases 12, 29 (2024).

3. Bennett, R. S. et al. Scalable, Micro-Neutralization Assay for Assessment of SARS-CoV-2 (COVID-19) Virus-Neutralizing Antibodies in Human Clinical Samples. Viruses 13, 893 (2021).

4. Frische, A. et al. Optimization and evaluation of a live virus SARS-CoV-2 neutralization assay. PLoS One 17, e0272298 (2022).

5. Cai, Z. et al. A Pseudovirus-Based Neutralization Assay for SARS-CoV-2 Variants: A Rapid, Cost-Effective, BSL-2–Based High-Throughput Assay Useful for Vaccine Immunogenicity Evaluation. Microorganisms 12, 501 (2024).

6. Münsterkötter, L. et al. Comparison of the Anti-SARS-CoV-2 Surrogate Neutralization Assays by TECOmedical and DiaPROPH-Med with Samples from Vaccinated and Infected Individuals. Viruses 14, 315 (2022).

7. Liu, K.-T., Han, Y.-J., Wu, G.-H., Huang, K.-Y. A. & Huang, P.-N. Overview of Neutralization Assays and International Standard for Detecting SARS-CoV-2 Neutralizing Antibody. Viruses 14, 1560 (2022).

8. Eliadis, P. et al. Novel Competitive ELISA Utilizing Trimeric Spike Protein of SARS-CoV-2, Could Identify More Than RBD-RBM Specific Neutralizing Antibodies in Hybrid Sera. Vaccines (Basel) 12, 914 (2024).

9. AVNIR technology for advanced neutralization assays. https://ivanobioscience.com/neutralization-assay-technology/.

10. Fiedler, S. et al. Antibody Affinity Governs the Inhibition of SARS-CoV-2 Spike/ACE2 Binding in Patient Serum. ACS Infect Dis 7, 2362–2369 (2021).

11. Klüpfel, J. et al. Automated detection of neutralizing SARS-CoV-2 antibodies in minutes using a competitive chemiluminescence immunoassay. Anal Bioanal Chem 415, 391–404 (2023).

12. Junker, D. et al. COVID-19 patient serum less potently inhibits ACE2-RBD binding for various SARS-CoV-2 RBD mutants. Sci Rep 12, 7168 (2022).

13. Najjar, D. et al. A lab-on-a-chip for the concurrent electrochemical detection of SARS-CoV-2 RNA and anti-SARS-CoV-2 antibodies in saliva and plasma. Nat Biomed Eng 6, 968–978 (2022).

14. Zhang, Y. et al. Development of receptor binding domain-based double-antigen sandwich lateral flow immunoassay for the detection and evaluation of SARS-CoV-2 neutralizing antibody in clinical sera samples compared with the conventional virus neutralization test. Talanta 255, 124200 (2023).

15. Lake, D. F. et al. Development of a rapid point-of-care test that measures neutralizing antibodies to SARS-CoV-2. Journal of Clinical Virology 145, 105024 (2021).

16. Liu, Z. et al. Development of an Effective Neutralizing Antibody Assay for SARS-CoV-2 Diagnosis. Int J Nanomedicine Volume 18, 3125–3139 (2023).

17. Bian, L. et al. Ultrabright nanoparticle-labeled lateral flow immunoassay for detection of anti-SARS-CoV-2 neutralizing antibodies in human serum. Biomaterials 288, 121694 (2022).

18. Mahmud, Md. A., et al. Semiquantitative Paper-Based Microfluidic Surrogate Virus Neutralization Test for SARS-CoV-2 Neutralizing Antibodies. Anal Chem 96, 11751–11759 (2024).

19. Rajil, N. et al. Quantum optical immunoassay: upconversion nanoparticle-based neutralizing assay for COVID-19. Sci Rep 12, 1263 (2022).

20. Cheedarla, N. et al. Rapid, high throughput, automated detection of SARS-CoV-2 neutralizing antibodies against Wuhan-WT, delta and omicron BA1, BA2 spike trimers. iScience 26, 108256 (2023).

21. Heggestad, J. T. et al. Rapid test to assess the escape of SARS-CoV-2 variants of concern. Sci Adv 7, (2021).

22. Bae, H. et al. Quantitative microfluidic assay to measure neutralizing and total antibodies for SARS-CoV-2. Sens Actuators B Chem 403, 135093 (2024).

23. Li, R. et al. Metasurface-Driven and Nanomaterial-Coupled Plasmonic Biosensor for the Rapid and Quantitative Clinical Identification of Neutralizing Antibodies against SARS-CoV-2 Variants. Adv Funct Mater 33, (2023).

24. Egia-Mendikute, L. et al. A flow cytometry-based neutralization assay for simultaneous evaluation of blocking antibodies against SARS-CoV-2 variants. Front Immunol 13, (2022).

25. Rahmati, Z., Roushani, M., Hosseini, H. & Choobin, H. An electrochemical immunosensor using SARS-CoV-2 spike protein-nickel hydroxide nanoparticles bio-conjugate modified SPCE for ultrasensitive detection of SARS-CoV-2 antibodies. Microchemical Journal 170, 106718 (2021).

26. Manshadi, M. K. D., Mansoorifar, A., Chiao, J.-C. & Beskok, A. Impedance-Based Neutralizing Antibody Detection Biosensor with Application in SARS-CoV-2 Infection. Anal Chem (2023).

